# Connectivity-Informed Adaptive Regularization for Generalized Outcomes

**DOI:** 10.1101/322420

**Authors:** Damian Brzyski, Marta Karas, Beau Ances, Mario Dzemidzic, Joaquin Goni, Timothy W Randolph, Jaroslaw Harezlak

**Affiliations:** Department of Epidemiology and Biostatistics, Indiana University, Bloomington, IN, USA; Johns Hopkins Bloomberg School of Public Health, Baltimore, MD, USA; Indiana University School of Medicine, Indianapolis, IN, USA; Purdue University, West Lafayette, IN, USA; Fred Hutchinson Cancer Research Center, Seattle, WA, USA

**Keywords:** Generalized Linear Regression, Penalized regression, Structured penalties, Laplacian matrix, Brain connectivity, Brain structure

## Abstract

One of the challenging problems in the brain imaging research is a principled incorporation of information from different imaging modalities in association studies. Frequently, data from each modality is analyzed separately using, for instance, dimensionality reduction techniques, which result in a loss of mutual information. We propose a novel regularization method, griPEER (generalized ridgified Partially Empirical Eigenvectors for Regression) to estimate the association between the brain structure features and a scalar outcome within the generalized linear regression framework. griPEER provides a principled approach to use external information from the structural brain connectivity to improve the regression coefficient estimation. Our proposal incorporates a penalty term, derived from the structural connectivity Laplacian matrix, in the penalized generalized linear regression. We address both theoretical and computational issues and show that our method is robust to the incomplete information about the structural brain connectivity. We also provide a significance testing procedure for performing inference on the estimated coefficients in this model. griPEER is evaluated in extensive simulation studies and it is applied in classification of the HIV+ and HIV- individuals.

## 1. Introduction

In brain imaging applications researchers often collect multiple data types, but in the majority of cases the analysis is performed separately for each of them. Implicit in the work of Randolph et al. (2012) is a framework for simultaneously utilizing multiple data types. For instance, structural and/or functional connectivity measures may serve as useful prior knowledge regarding the structure of dependencies between brain regions when used in a linear model that aims to estimate the association of brain region properties (e.g., cortical thickness) with a scaler outcome. Karas et al. (2017) explicitly showed that using correct prior information significantly increases estimation accuracy. The statistical methodology, riPEER, developed by these authors allows for incorporating such predefined structure into a regression model a way that protects against using incorrect information. The derived estimation procedure, however, is limited by the assumption that the response variable is normally distributed. Such design excludes, for instance, a binary response that indicates the presence/absence of a condition such disease or phenotype.

To fill this gap, we developed a variant of riPEER, called *generalized ridgified Partially Empirical Eigenvectors for Regression* (griPEER), which handles the outcomes coming from the exponential family of distributions. In the context of brain imaging analysis, our approach allows the analysis to incorporate information such as that encoded in a structural or functional connectivity matrix. As with its precursor, griPEER is able to use the predefined information in a “soft” way — from full inclusion absence — depending on how well this information is confirmed by the data. To achieve this, griPEER employs a penalized optimization problem with a flexible, parameterized penalty term with parameters chosen in a fully automatic and data-driven manner.

We work with a generalized linear regression model where the *i*th scalar outcome, *y*_*i*_, is assumed to be drawn from the exponential family of distribution with the parameter *θ*_*i*_. We confine ourselves to the canonical link functions only and assume that *θ* = *Xβ* + *Zb*. Here, *X* denotes a matrix of covariates (such as demographic data) for which the prior information is not used and the columns of *Z* correspond to variables having structure which is assumed to be at least partially known. In the analysis performed in this article, *β* includes the intercept and demographic data, while *b* represents the coefficients for the average thickness of 66 brain regions. These regions are assumed to be linked and this linkage is represented by a connectivity matrix; e.g., this matrix may encode a density of connections or the average Fractional Anisotropy (FA).

There is a wide literature on using structural information in image reconstruction and estimation (see, e.g., Bertero and Boccacci (1998), Engl et al. (2000), Phillips (1962)). In situations when the object of the interest is assumed to be a function belonging to a class of, say, differentiable functions, a differential operator-based penalty may be used to “regularize” or impose smoothness on the estimates (Huang et al., 2008). This may improve the prediction and interpretability and is “efficient and sometimes essential” in situations having many highly correlated predictors (Hastie et al., 1995). When the object of estimation is a vector, the penalties are very often constructed based on ℓ_1_ and ℓ_2_ norms. Examples include such methods as LASSO (Tibshirani, 1996), adaptive LASSO (Zou, 2006), ridge regression (Tikhonov, 1963) and elastic net (Zou and Hastie, 2005), to name just a few.

There is no the unique answer to the question of how to regularize a particular model and the final construction depends strongly on the context. If, for instance, it is natural to assume sparsity in the coefficients or that they occur in blocks, then using the ℓ_1_ norm to constrain them (as in the LASSO) or constrain the difference of adjacent coefficients (as in the fused LASSO) would be useful (Tibshirani et al., 2005). A more generalized fused lasso using two ℓ_1_ norms could also be applied: one constraining on the coefficients and one constraining their pairwise differences (Xin et al., 2016).

When more intricate structure among the variables is expected and when some (possibly imprecise) knowledge of it is available, then less generic penalization schemes are more appropriate (Tibshirani and Taylor, 2011; Slawski et al., 2010). For example, a *p* × *p* adjacency matrix represents known connections, or “edges”, between *p* nodes in a graph. This matrix can be used to inform a model that aims to estimate the relationship between an outcome an a vector of *p* values at the nodes in the graph. More specifically, the adjacency matrix is used to define the graph Laplacian matrix which represents differences between nodes (Chung, 2005), and may be used to penalize the process of estimating regression coefficients, *b*.

For any *p* × *p* matrix *Q*, defining a penalty of the form λ*b*^T^*Qb*, where λ is a nonnegative regularization parameter constitutes the essence of the methods of Li and Li (2008) and Karas et al. (2017). Using a penalty of this form also serves to link the optimization problem with theory of mixed effects models in which *b* is assumed to be a random effects vector with distribution 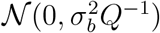, for some 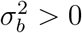. This, in turn, reveals a connection with the Bayesian approach, where the distribution is treated as a prior on *b*; see e.g., Maldonado (2009).

Problems with such an interpretation include the fact that *Q* may not be invertible, as is the case when *Q* is defined as Laplacian or normalized Laplacian (Chung, 2005). Second, a single multiplicative parameter, λ, adjusts the trade-off between model fit and penalty terms but it can not change the regularization pattern, i.e., the shape of the set {*b* : *penalty*(*b*) = *const*} is preserved. When *Q* is misspecified (is not informative) this lack of adaptivity may significantly degrade performance to be even worse than a uninformed penalty such as ridge regression or LASSO (Karas et al., 2017).

Both of these issues were considered in (Karas et al., 2017) which does not assume *Q* is exactly the true signal precision matrix, ℚ, but is merely“close”, in some sense; i. e., *Q* contains some amount of true information which can be exploited. Therefore, by considering a family of transformations of *Q*, and selecting the optimal member by applying a data-adaptive procedure, one may obtain a modified matrix which reflects ℚ better and improves prediction accuracy. Transformations of the form *Q* + *a***I**_*p*_ (*a* > 0) are used by Karas et al. (2017). Any such modification of *Q* is invertible and could be directly used in the estimation procedure. The resulting penalty term, λ*b*^T^(*Q* + *a***I**_*p*_)*b*, has an equivalent form 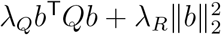 and the connection with a specific linear mixed model enables the optimal selection of λ_*Q*_ and λ_*R*_.

The approach by Karas et al. (2017) assumes the response variable is normally distributed and hence not suitable for categorical outcomes. In this presentation we extenbd the concept of riPEER’s penalty function to the case when the distribution of the response variable is a member of one-parameter exponential family of distributions. The proposed estimation method, griPEER, is of the form:

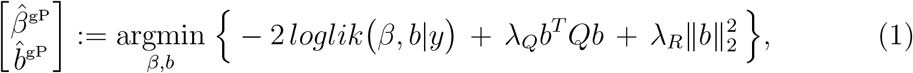

where *loglik*(*β*, *b*|*y*) is a log-likelihood. Here, the term ‒*2loglik*(*β*, *b*|*y*) is used to fit the model to the response distribution while the parameters λ_*Q*_ and λ_*R*_ are chosen based on the connection between the optimization problem and the generalized linear mixed model; this is formulated explicitly in Section 2. It is important to emphasize that these parameters not only determine the trade-off between the model fit and the penalty term, but also on the form of the penalty, which determines the structure that the estimate is encouraged to have. More precisely, if λ_*Q*_ is large relative to λ_*R*_, then the connectivity information has a large role in the estimation process. Conversely, when λ_*Q*_ is small relative to λ_*R*_, the penalty is equal in all coordinates, as with ridge regression.

We illustrate this using a simple example with *p* = 2 variables and prior information that implies these variables are connected. Figure 1 shows how the shapes of contour sets of penalty, which decide on the solution structure, change for various lambdas. If the relationship between variables, as represented in *Q*, is reflected in data and if this is related to the outcome *y*, then griPEER will tend to choose relatively large λ_*Q*_, which links the coefficients in *b* (see right plot in Figure 1). The other extreme is when the structure in *Q* is not informative for the relationship between *y* and *Z*. In this case, griPEER will select a relatively large λ_*R*_ inducing a ridge-like penalty that ignores *Q* (see the left panel in Figure 1).

**Figure 1:**
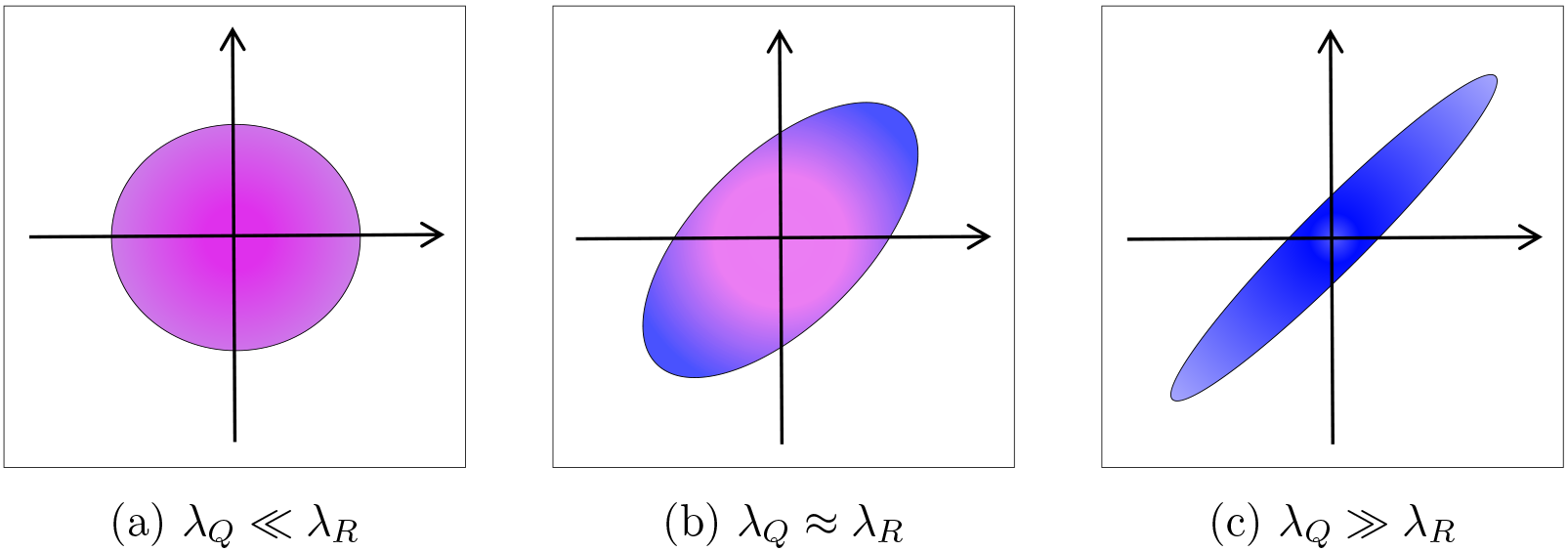
Shapes of the set 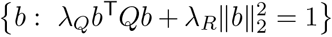 for various pairs of regularization parameters: (a) the assumed strong connections between variables was neglected, (b) the moderate tendency for coefficients of the solution to be similar to each other (c) strong tendency for coefficients of the solution to be similar to each other

The remainder of this work is organized as follows. In Section 2, we formulate our statistical model, investigate the special case of binomial distribution and discuss the equivalence between GLMM and penalized optimization problems. We also describe the penalty term construction from the graph-theory point of view. The estimation procedure we use to select the optimal regularization parameters is introduced in Section 3, while the next Section addresses the problem of the selection of response-relevant variables. The extensive simulations showing very good performance of griPEER in the context of estimation accuracy and variables selection (under various scenarios illustrating the impact of inaccurate prior information) are reported in Section 5. Finally, in Section 6, we apply our methodology to study the association of the brain’s cortical thickness and HIV disease. The conclusions and a discussion are summarized in Section 7.

## 2. Statistical model

We address the problem of estimation in a penalized generalized linear model where the penalty term is derived from connectivity information. This information is represented by a *p* × *p* symmetric matrix having non-negative entries with zeros on the diagonal. This *adjacency matrix* or *connectivity matrix* and will be denoted by ***A***. The corresponding graph Laplacian matrix, *Q*, which defines the penalty term is defined next, followed by specific details about the statistical model in (1).

### 2.1. The graph Laplacian, Q

We are interested in modeling the association between a scaler outcome, *y*, and a set of *p* predictor variables that are measured at the nodes of graph. We assume that information about connections between the these variables — i.e., strengths of the connections between the nodes — can be summarized by a (symmetric) *p* × *p* adjacency matrix 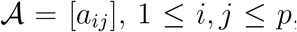, having non-negative entries and zeros on the diagonal. We denote the *degree* of the *j*th node as *d*_*j*_ := ∑_*j*_ *a*_*jj*_, and define the degree matrix as *D* := *diag*(*d*_1_,…, *d*_*p*_).

Following Chung (2005), we define the unnormalized Laplacian, *Q*_*u*_, corresponding to 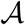 simply as 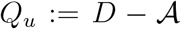. This matrix is always positive semidefinite. It is also singular, since for the vector of ones, **1** := [1,…, 1]^T^ we have 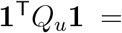 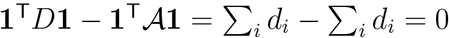.

Intuition on the role of a penalty of the form *b*^*T*^*Q*_*u*_*b*, as in (1), is gained by the following simple formula: for any adjacency matrix, 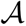, and its unnormalized Laplacian, *Q*_*u*_, then

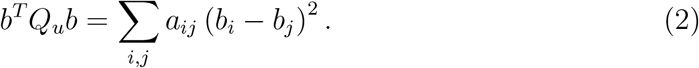

That is, the term *b*^*T*^*Q*_*u*_*b* in the optimization problem (1) penalizes the squared differences of coefficients in a manner that is proportional to the strengths of connections between them. Consequently, coefficients corresponding to nodes having many strong connections (nodes with large degree) are constrained more than others. T

In order to allow a small number of nodes with large *d*_*i*_ to have more extreme values, we employ the normalized Laplacian, *Q*, which is obtained by by dividing each column and row of *Q*_*u*_ by a square root of corresponding node’s degree. As a result, the property (2), with *Q* instead of *Q*_*u*_, takes the form

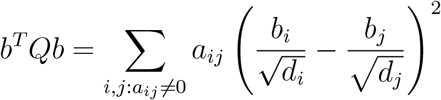

*Q* has ones one the diagonal and, as with the unnormalized Laplacian, it is a symmetric, positive semidefinite and singular matrix.

### 2.2. Statistical model in general form

Consider the general setting where *y* is an *n* × 1 vector of observations, and the design matrices, *X* and *Z*, are *n* × *p* and *n* × *m* matrices, respectively. The columns of *X* represent the *p* covariates and the rows are denoted by *X*_*i*_. Similarly, the columns of *Z* correspond to *m* variables, or nodes in a graph, for which some connectivity information may be available; the rows are denoted by *Z*_*i*_. We assume there exists(unknown) vectors *b* and *β* such that, for each *i* ∈ {1,…, *n*}, *y*_*i*_ is the member of one-parameter exponential family of distributions of the form

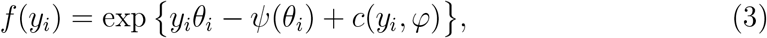

where *θ*_*i*_:= *X*_*i*_*β* + *Z*_*i*_*b* is a subject-specific parameter. The formula in 3 includes exponential, binomial, Poisson and Laplace densities.

It can be shown that for the exponential family of distributions, the mean of *y*_*i*_ is simply given by the first derivative of *ψ* in the point *θ*_*i*_, while the variance could be expressed as the second derivative of *ψ*, i. e.

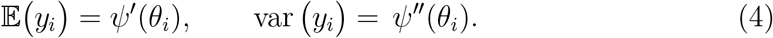

Moreover, the log-likelihood function is

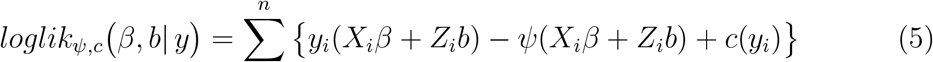

and it provides a core for the methodology we propose in this presentation. Indeed, we define griPEER as a solution to the following optimization problem

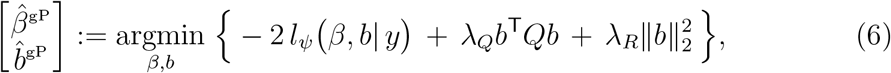

where 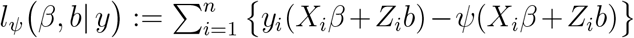 consists of the terms of log-likelihood function (5) depending ond *b* and *β*. Here, λ_*Q*_ and λ_*R*_ are regularization parameters, which are selected automatically, as described in Section 3.

### 2.3. The special case – binomial distribution

To provide focus to our presentation we will concentrate on the setting of a binomial outcome in all simulations (Section 5) and the responses in the applications (Section 6) are modeled by the binomial distribution. So in this subsection we explicitly describe this special choice of density function.

As in the classical logistic regression theory, we assume that the response, *y*_*i*_, takes the value 1 with probability 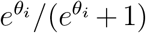 and 0 with the probability 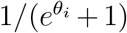. Consequently, the density function, *f*(*y*_*i*_), is given by

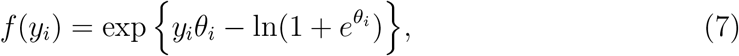

which is a member of exponential family of distributions (3) with 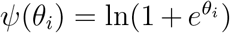 and *c*(*y*_*i*_) = 0. We also have

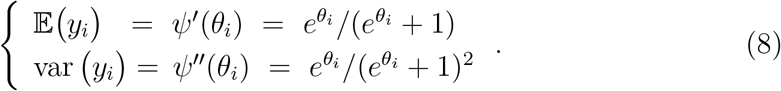

From this, 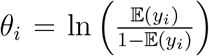 which, with the assumption *θ* = *Xβ* + *Zb* adopted in
the manuscript, yields the canonical link for logistic regression—the logit function.

### 2.4. Equivalence between GLMM and two optimization problems

The optimization problem in (6) is strongly connected with the specific GLMM formulation. Indeed, consider the model defined by the following conditions

**A.1** *β* is a vector of fixed and *b* is a vector of random effects,
**A.2** *y*_*i*_|*b* are independent and, consequently, 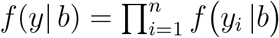,
**A.3** *f*(*y*_*i*_ |*b*) = exp {*y*_*i*_(*X*_*i*_*β* + *Z*_*i*_*b*) − *ψ*(*X*_*i*_*β* + *Z*_*i*_*b*) + *c*(*y*_*i*_)}, for some (known) functions *ψ*, *c* and *i* = 1,…,*n*,
**A.4** 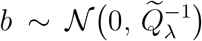, where 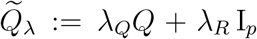 for some unknown, positive parameters λ_*Q*_ and λ_*R*_.

To see this correspondence, assume the parameters λ_*Q*_ and λ_*R*_ have been esti-mated, say as 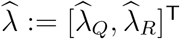, and these values are used to obtain *β* and *b*. One can proceed by treating both fixed and random effects as parameters and finding ML estimates by maximizing (with respect to *β*, *b*) the density function

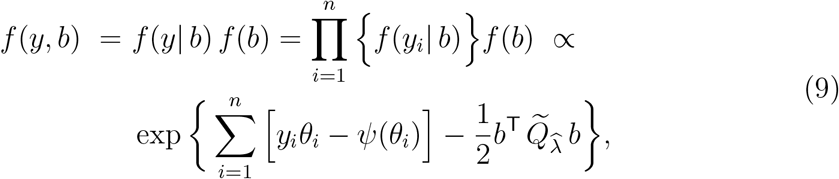

where *θ*_*i*_ = *X*_*i*_*β* + *Z*_*i*_*b*, for *i* = 1,…,*n*. Taking the logarithm of the above leads directly to the objective in optimization problem (6).

We now derive a constrained optimization problem that is equivalent to (6) and reveals the role of the regularization parameters on the solution from a slightly different perspective. For this, suppose that 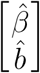 is the solution to (6) for given parameters λ_*Q*_ and λ_*R*_. Then define 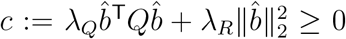. One can check that 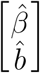 also solves the problem

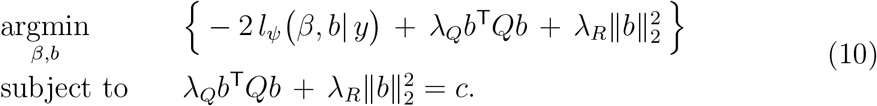

The multiplicative factor may be neglected as well as the term 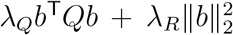, which is constant on the feasible set. This yields

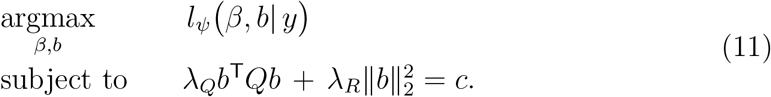

This formulation clarifies the intuition behind the example in the Introduction and the corresponding Figure 1. I.e., griPEER selects the estimates by taking the maximal likelihood value on a set whose shape is explicitly regularized by the parameters λ_*Q*_ and λ_*R*_.

## 3. A new estimation algorithm

To select the optimal values of λ_*Q*_ and λ_*R*_, we employ the corresponding GLMM formulation defined by A.1 – A.4. The likelihood function, 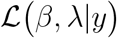, is given by

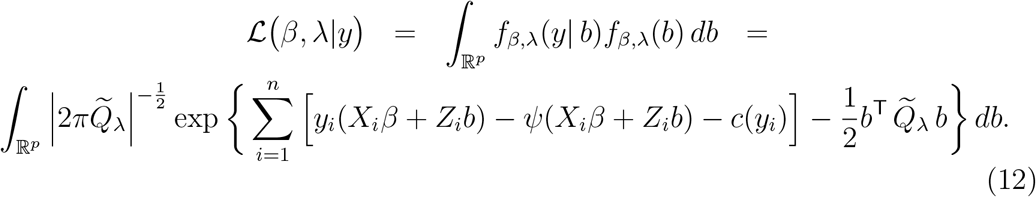

Unfortunately, obtaining the maximum of 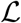 with respect to *β* and λ is complicated by the fact that there is no closed-form solution to the multidimensional integral in (12). For this, several approaches have been proposed. Breslow and Clayton (1993) proposed a general method based on Penalised Quasi-Likelihood (PQL) for the estimation of the fixed and prediction of random effects. Wolfinger and O’connell (1993) investigated the pseudo-likelihood (PL) approach which is closely related to the Laplace’s approximation of 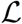. Other proposals include the Adaptive Gaussian Quadrature to approximate integrals with respect to a given kernel (Pinheiro and Chao, 2006) and an MCMC-based procedure (Zeger and Karim, 1991).

In this article we focus on the Wolfinger PL approach which is recognized as being fast and computationally efficient. It relies on the first-order Taylor series approximation and uses the Linear Mixed Model (LMM) proxy in the iterative process: at each iteration, the updates of *β* and *b* are based on the variance-covariance parameters of random effects. The steps are repeated until convergence.

The procedure we derive here differs from (Wolfinger and O’connell, 1993) in how the updates of *β* and *b* are obtained. In contrast to the Wolfinger PL approach, we do not get them via the solution to the mixed-model equations, but instead we employ the correspondence between GLMM and griPEER optimization problem, as described in 2.4. Specfically, the (*k* – 1)-step estimates of λ_*Q*_ and λ_*R*_ (i.e., 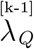 and 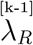) are used to obtain the (*k* - 1)-step estimates of *β* and *b* (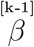 and 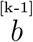) via the solution to (6). Consequently, we can define 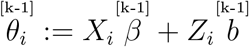.

Details of our estimation procedure are as follows. Using the Taylor approxima-tion of function *ψ*′ at point 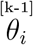 we get

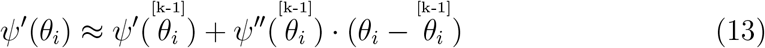

and therefore from (4)

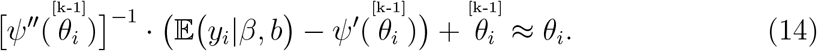

We now define a random variable 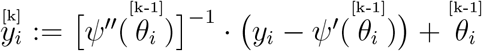. The main step now is the assumption that the distribution of 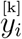 can be well approximated by a normal density. Computation of mean and variance of 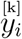 immediately yields

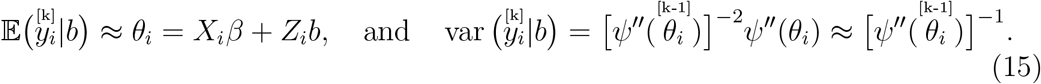

The assumption that 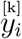 is approximately normally distributed allows for replacing the GLMM formulation in *k*th step by an LMM of the form

**B.1** *β* is a vector of fixed and *b* is a vector of random effects,
**B.2** 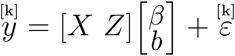,
**B.3** 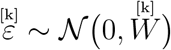, where 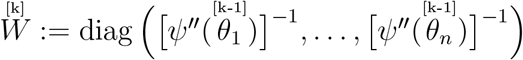,
B.4 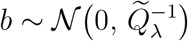, where 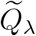 was defined in (A.4).

Denote by 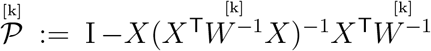 the 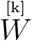-weighted projection onto the orthogonal complement of the columns of *X*. Now, defining 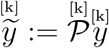,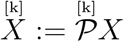 and 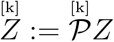 we assume that

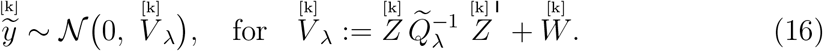

Maximizing the log-likelihood for 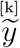, i.e. the function 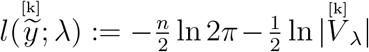 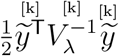, leads directly to the optimization problem

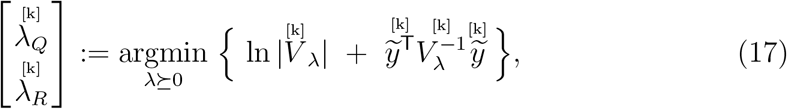

where λ ⪰ 0 refers to {(λ_*Q*_, λ_*R*_) : λ_*Q*_ ≥ 0, λ_*R*_ ≥ 0}. The following proposition helps us to rewrite the objective of (17). A proof is provided in the Appendix.

### Proposition 3.1

*Let* 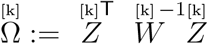 *and* 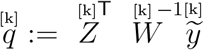. *Then*

***C.1*** 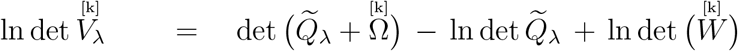,
***C.2*** 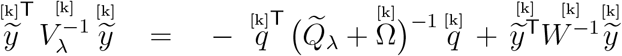.

This proposition makes it possible to reformulate (17) and define the *k*th step update, 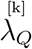 and 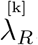, as

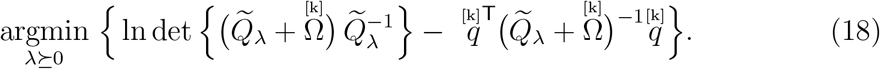

It is important to use an efficient and accurate method to solve (18) since this problem appears in every step *k* and determines when the entire algorithm terminates (when 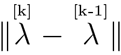 is sufficiently small). To achieve this, we have analytically derived the gradient and the Hessian of the objective function. (Details are in the the Appendix.) The final algorithm for selecting the regularization parameters is outlined here:

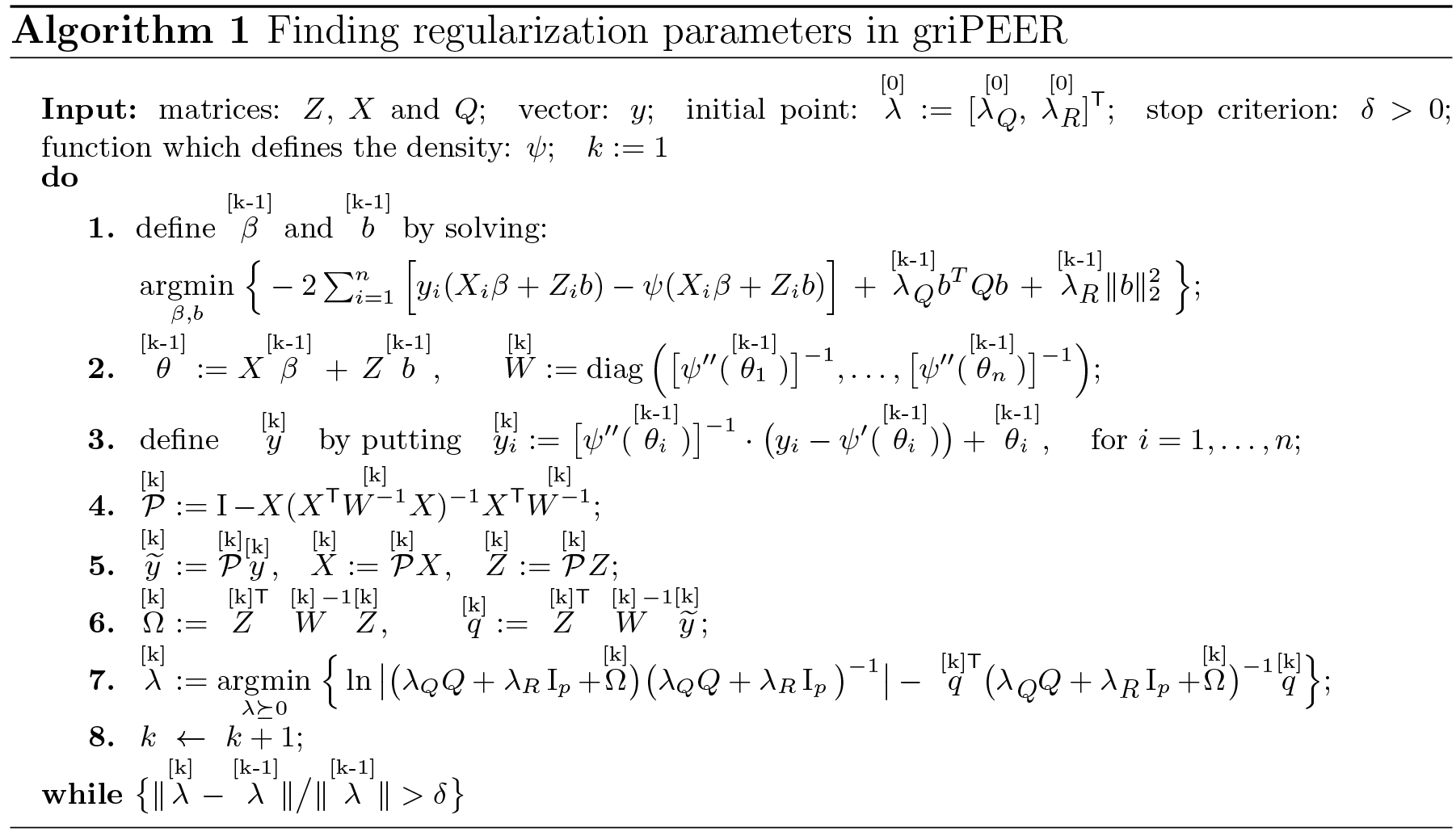

## 4. Procedures for the significance testing

Unlike the lasso estimation procedure that produces a sparse set of regression coefficients but does not (without additional theory Zhao and Shojaie (2016)) provide statistical significance testing, we employ two methods to identify variables that are identified as statistically significantly related to the response. Two such approaches are implemented in our software and we introduce them in this section. They both use the knowledge about the optimal regularization parameters described in the previous section. The first takes advantage of asymptotic properties of generalized linear model (GLM) estimates and construct the estimate of asymptotic variance-covariance matrix in the similar fashion as proposed by Cessie and Houwelingen (1992) in the context of ridge-penalized logistic regression. The second applies the bootstrap method. When griPEER is used for variable selection, we will refer to these two approaches as griPEER_asmp_ (the asymptotic-based approach) and griPEER_boot_ (the bootstrap-based approach), respectively. The numerical experiments performed in Section 5 suggest that griPEER_boot_ is able to achieve significantly larger power than griPEER_asmp_ under the settings reflecting brain imaging design and connectivity matrices. Since the same experiment shows similar rates of false discoveries among variables labeled as relevant, griPEER_boot_ was used in real data analysis (Section 6) to find brain regions associated with HIV.

### 4.1. Asymptotic variance-covariance matrix

We start by introducing notation. Denote 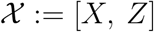, let 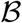 be *p*+*m* dimensional estimate given by (6), and 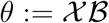. Moreover, we define a (*p*+*m*)×(*p*+*m*) penalty matrix as

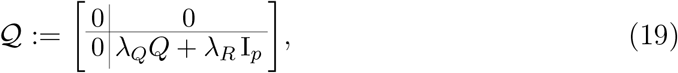

where the non-negative parameters λ_*Q*_, λ_*R*_ are adjusted by the procedure 1. In summary, 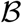 is the solution to

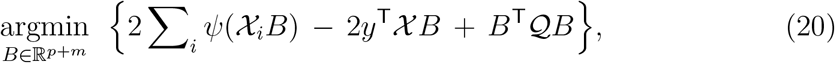

with *ψ* being a given function indicating the member of the exponential family of distributions (3). Furthermore, the formulas we derive in this section include the diagonal matrix Ψ defined as Ψ := diag {*ψ*″(*θ*_1_),…,*ψ*″(*θ*_*n*_)}.

Using the first-order Taylor approximation, as well as asymptotic properties of GLM estimate, one can find that the estimate asymptotic variance for 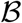 has a form

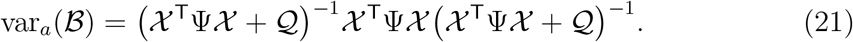

The derivation is based on Cessie and Houwelingen (1992) and was described in detail in Appendix A.3. Based on the above formula, we propose a simple decisionmaking strategy in which we label the *i*th covariate as statistically relevant if 0 is not included in the 95% confidence interval for its respective regression coefficient,
i.e.

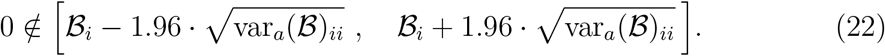

### 4.2. The Bootstrap based approach

In this approach the variances of coefficients in 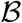, the solution to (20), was estimated based on Bootstrap samples. Each such sample was created from *n* elements of *y* and *n* corresponding rows of *Z* and *X*, which indices were selected randomly by sampling with replacement. The dataset obtained in *j*th repetition, *X*^[*j*]^, *Z*^[*j*]^ and *y*^[*j*]^, were then substituted to the objective in (6) with λ_*Q*_ and λ_*R*_ being selected by Algorithm 1 applied to the original dataset (i.e., λ_*Q*_ and λ_*R*_ were estimated only once). The percentile bootstrap confidence intervals, with the significance level *α* = 0.05, were defined based on all estimates, ***B***^[1]^, …, ***B***^[*s*]^. The default value of *s* was set to 500 in our software and this number of bootstrap samples was generated in simulations performed in subsection 5.3. Coefficients from the griPEER estimate whose confidence intervals do not contain zero are labeled as response-related discoveries.

## 5. Numerical experiments

We conduct a simulation study to investigate the performance of griPEER in the situation when responses are modeled by binomial distribution. Results are compared with the logistic ridge estimates.

### 5.1. Definitions

*Matrix density*. For a *p* × *q* matrix *A* define its *density* as a proportion of non-zero entries,

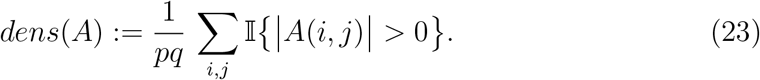

*Matrix dissimilarity*. To quantify a dissimilarity between two *p* × *q* matrices, *A* and *B*, with *dens* (*A*) = *dens*(*B*), we define

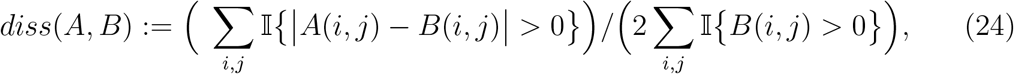

with values in the interval [0,1]. If *diss*(*A*,*B*) = 0 then *A* = *B* and *diss*(*A*,*B*) = 1 means that the positions of non-zero entries do not overlap.

### 5.2. Model coefficient estimation

#### 5.2.1. Settings

*“Informativeness” of the penalty term*. The simulation settings were designed to evaluate performance in a variety of situations ranging from an “observed” connectivity matrix (i.e., a prescribed matrix used in estimation) that is fully informative to one that is completely non-informative. Here “informativeness” refers to the amount of true dependencies among the variables that are represented in the connectivity matrix.

Denote By 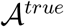 a matrix representing true connections between variables and by 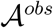 one which is observed and used in an estimation via griPEER. To express “informativeness” of 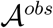 with respect to 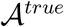, we use a measure of dissimilarity, 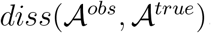, defined in (24). We have

- 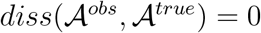 reflects a situation when 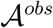 is fully informative;
- 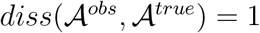 reflects a situation when 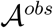 is non-informative;
- 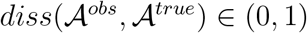 indicates 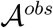 is partially informative.

*Connectivity in the context of brain regions*. One may view 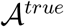 as an adjacency matrix of a graph representing the connections between brain regions, and our simulations scenarios are based on the following four interpretations regarding this structure.

1. 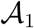: “*homologous regions*”. 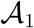 represents a situation when brain regions, *i* and *j*, are connected (i.e., 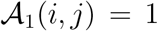) if *i* and *j* are homologous brain regions from different hemispheres, and 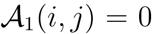 otherwise. This matrix is shown in Figure 2, left plot.
2. 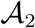: “*modularity*”. 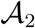 represents a situation when brain regions *i* and *j* are connected if and only if they belong to the same *module* with 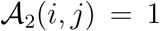 within the module and 0 otherwise. This matrix is shown in Figure 2, middle left plot.
3. 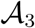: “*density of connections*, *masked*”. 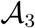 is defined based on the brain-imaging measure — *density of connections between brain regions* (see, Section 6) — and then is “masked” by modularity information. Here, 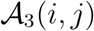 equals the median of a density of connections between regions *i* and *j* if they belong to the same module. Otherwise, 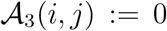. Matrix 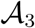 is shown in Figure 2, middle right plot.
4. 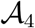: “*neighboring regions*”. 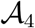 represents a situation when brain regions *i* and *j* are connected if they are “close” according to their spatial location 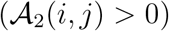. Otherwise, they are not connected 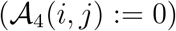. This matrix is shown in Figure 2, right plot.

**Figure 2:**
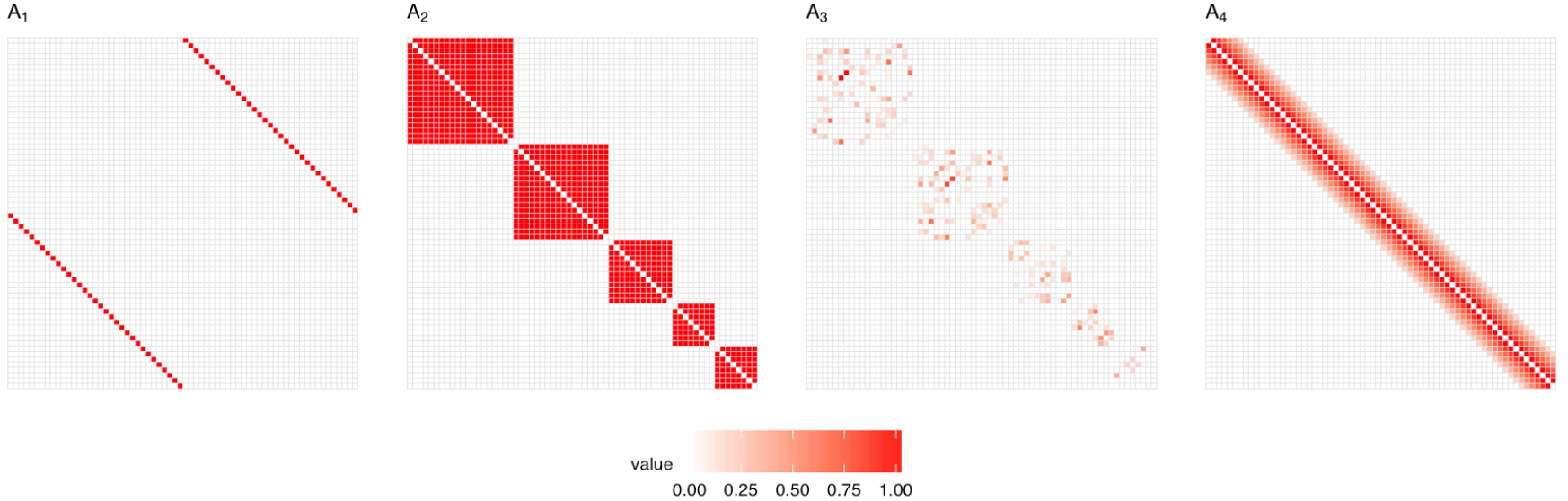
Matrices used in the simulation study to construct 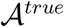. Presented are variants for *p* = 66. Left plot: 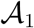 “*homologous regions*”. Middle left plot: 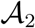 “*modularity*”. Middle right plot: 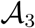 “*density of connections, masked*”. Right plot: 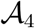 “*neighboring regions*”.

A *homologous regions* matrix 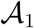 reflects the situation where only homologous regions from two hemispheres are assumed to be connected. A *modularity* matrix 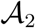, in turn, represents adjacency defining division of the brain cortical regions into five modules (Sporns, 2013; Cole et al., 2014; Sporns and Betzel, 2016). Next, a “*density of connections*, *masked*” matrix 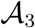 is based on estimated density of connections between brain cortical regions, as described in Section 6). Finally, the “*neighboring regions*” matrix 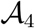 models the situation where brain regions are spatially connected; i. e., the strength of connection between brain regions depends on the physical distance between them.

*Simulation scenarios*. We run three simulation scenarios to express different sources of “uninformativeness” 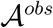 which loosely reflect real-life scenarios. For each scenario, we tested all four types of matrices, 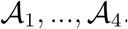.

- **Scenario 1.** The observed connectivity matrix, 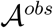, represents connections (partially) permuted with respect to connections represented by 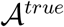. Based on one of four considered matrices, the corresponding 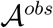 matrix is constructed by randomizing edges of a graph given by 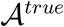 until a desired dissimilarity, 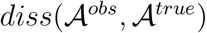, is achieved (see: Fig. 3). The randomization technique preserves graph size, density, strength and graph degree-sequence (and hence degree distribution).
- **Scenario 2.** We investigate the impact of using inaccurate information by labeling the negative connections between variables as the positive. For 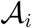, with *i* ∈ {1,…,4}, the true signal was generated after changing the structure of variables dependencies by allowing some negative connections first. Specifically, 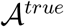 was defined by turning entries of columns *k* ∈ {1, 4, 7,10} and corresponding rows of 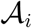 into their negative values. Here, 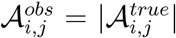 and hence 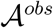 contains only non-negative values.
- **Scenario 3**. The observed connectivity matrix 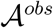 is of lower or higher matrix density than 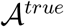. For 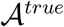 defined based on one of four considered matrices, the corresponding 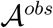 is constructed by randomly removing, respectively adding, edges to the graph of connections represented by 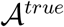 until the desired ratio of matrix densities, 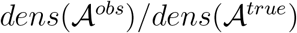, is obtained (see Fig. 5).

**Figure 3:**
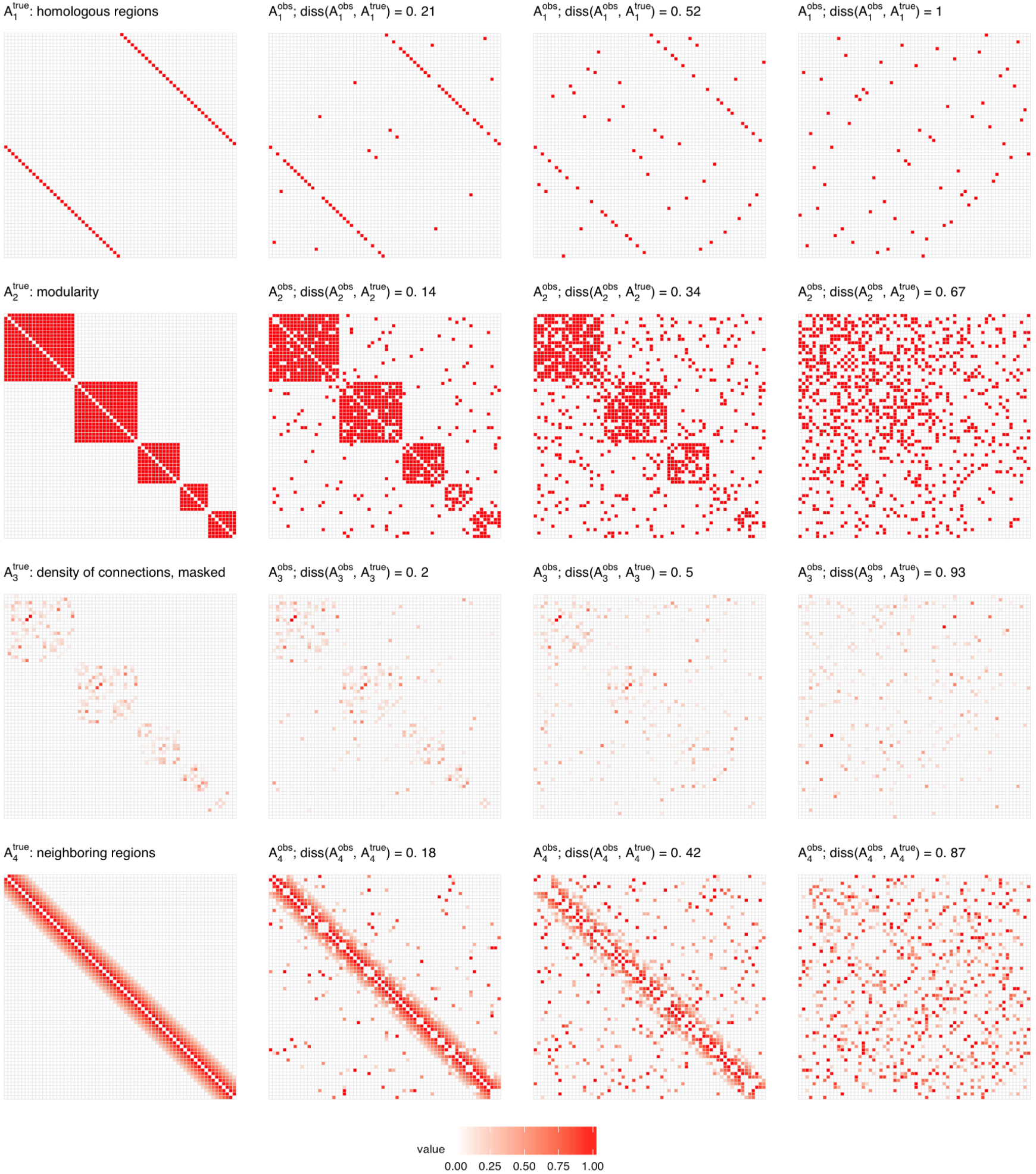
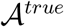 connectivity graph adjacency matrices (1st column panel) and 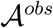 connectivity graph adjacency matrices (2nd-4th column panels) used in Scenario 1. 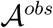 matrix is constructed by randomizing 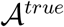 until a desired dissimilarity, 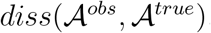, is achieved (*diss* is growing when moving from left to right side of each row plot panel).

*Simulation procedure*. In each numerical experiment, we perform the following steps.

1. For graph adjacency 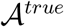, compute its normalized Laplacian, *Q*^*true*^ (in Scenario 2 the node’s degree is defined as *d*_*i*_ := ∑_*j*_ |*a*_*ij*_|; see subsection 2.1).
2. Replace the zero singular values of *Q*^*true*^ by 0.01 · *s*, where *s* is the smallest nonzero singular value of *Q*^*true*^ to get an invertible matrix required in 6. (a)).
3. For graph adjacency matrix, 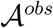, compute its normalized Laplacian, *Q*^*obs*^.
4. Generate *Z* ∈ ℝ^*n*×*p*^, where the rows are independently distributed by 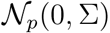, where ∑ is variance-covariance matrix estimated from a real data study (see: Sect. 6); standarize columns of *Z* so as they have mean 0 and unit ℓ_2_ norm.
5. Generate *X* as *n*-dimensional column of ones.
6. Run the following steps 100 times:

a. generate *b* ∈ ℝ^*p*^ as 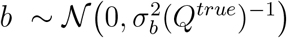; set *β* = 0,
b. define *θ* := *Xβ* + *Zb*,
c. define 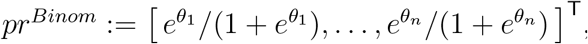,
d. generate *y* ~ *Binom*(*pr*^*Binom*^), *y* ∈ ℝ^*n*×1^,
e. estimate model coefficients *b*, *β* with the two methods: (1) griPEER, assuming the binomial distribution of *y* and using *Q*^*obs*^ in a penalty term, (2) logistic ridge estimator,
f. compute *b* estimation error, 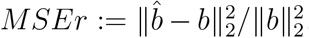,for two *b* estimates, (1) 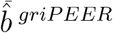 and (2) 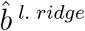.
7. Compute mean MSEr out of the 100 runs from (5), for the two estimation methods.

Importantly, a “true” coefficient vector *b* obtained as 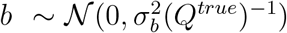 reflects the connectivity structure represented by 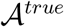. Examplary vectors *b* generated based on 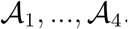 are presented in Figure 11 in Appendix B.

**Figure 4:**
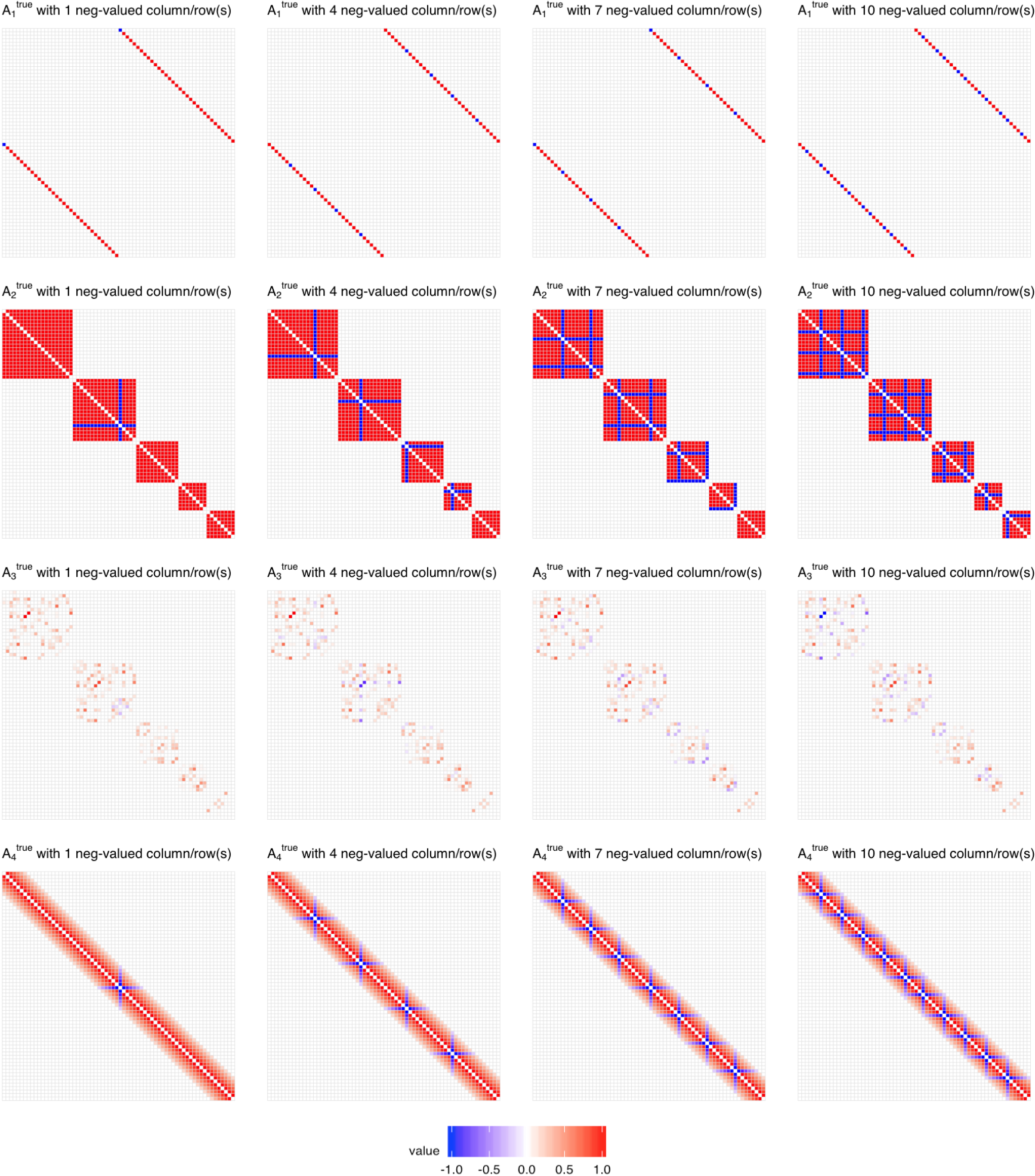
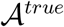 connectivity graph adjacency matrices used in Scenario 2. 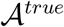 matrix is constructed from 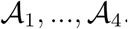 matrices (1st-4th row panels, respectively) by turning entries of *k*, *k* ∈ {1, 4, 7,10}, columns (and corresponding rows) of this matrix into their negative values (k is growing when moving from left to right side of each row plot panel).

**Figure 5:**
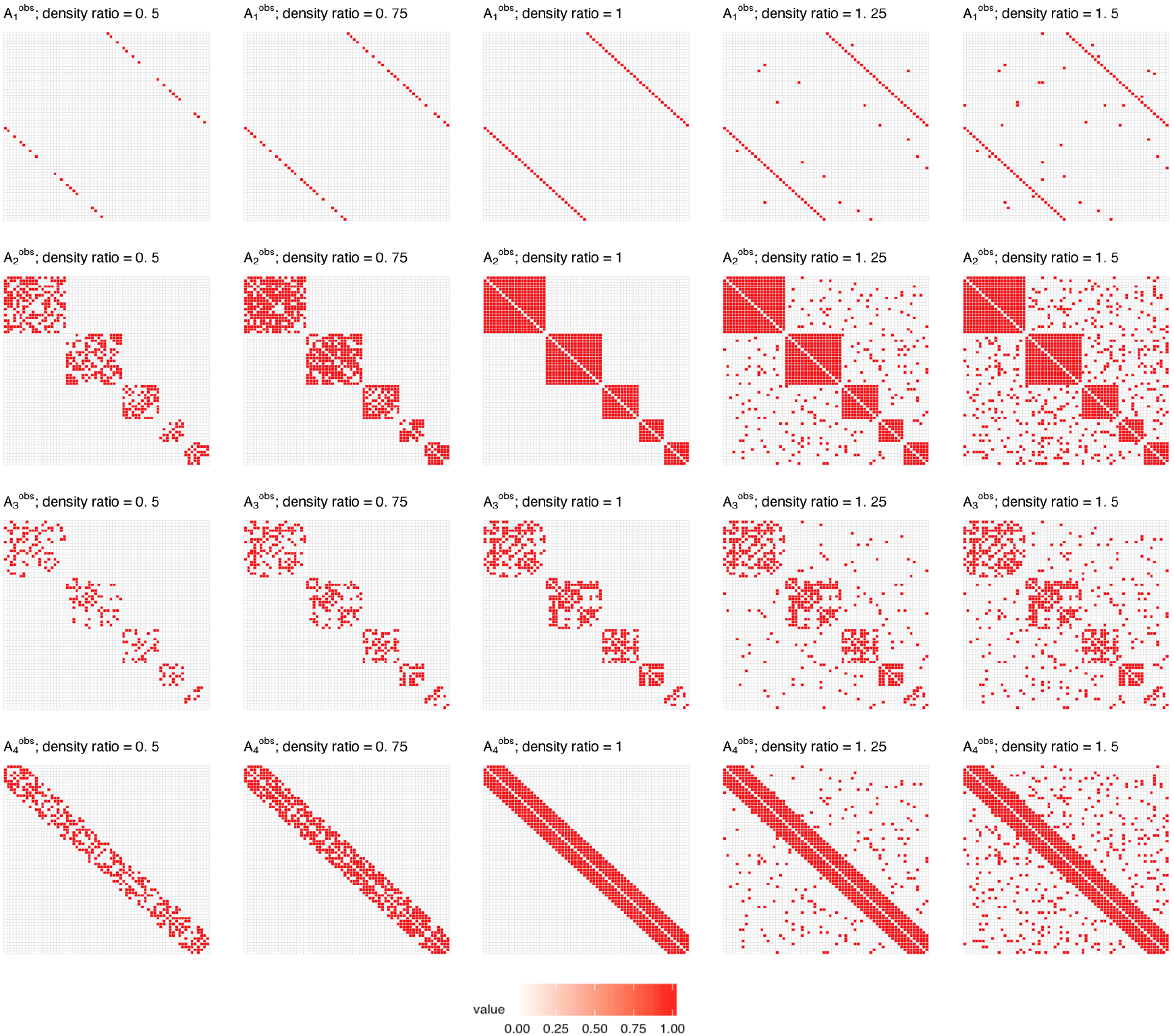
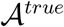 and 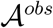 connectivity graph adjacency matrices used in Scenario 3. 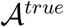 matrix is defined as one of 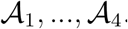 matrices (1st-4th row panels, respectively). Corresponding 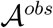 is constructed by randomly removing / adding edges to the graph of connections represented by 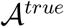 until desired density ratio, 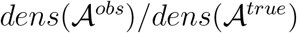, is obtained (ratio is growing from 0.5 to 1.5 when moving from left to right side of each row plot panel).

*Simulation parameters*. We consider the following choices of the experimental settings:

1. number of predictors: *p* ∈ {66,198},
2. number of observations: *n* ∈ {100, 200},
3. 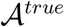 matrix constructed based on 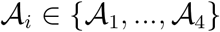,
4. (Scenario 1.) dissimilarity between 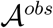 and 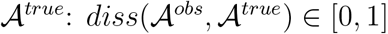,
5. (Scenario 2.) number of columns (and corresponding rows) of 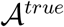 that have switched signs: *k* ∈ {0,1, 4, 7,10},
6. (Scenario 3.) density ratio: 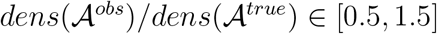.

The number of predictors, *p* = 66, is motivated by the brain imaging analysis described in Section 6, where 66 brain regions were considered. To investigate the situations with larger number of predictors for *i*th type of connectivity pattern, we created block-diagonal adjacency matrices with 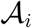’s as blocks. The adjacency matrix in the case with *p* = 198 was therefore defined as 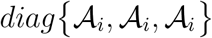.

#### 5.2.2. Results

*Scenario 1*. In Scenario 1, we compare griPEER and logistic ridge estimation methods in a situation when an observed connectivity matrix 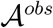 contains connections that are permuted with respect to connections represented by 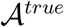. We consider combinations of simulation parameter values: number of predictors *p* ∈ {66,198}, number of observations *n* ∈ {100, 200}, 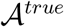 base matrix 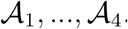, dissimilarity between 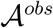 and 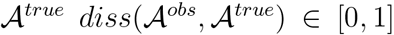. Fig. 6 displays the aggregated (mean) values of the relative estimation error based on 100 simulation runs.

We observe that in each case, MSEr of griPEER is lower or equal to MSEr of logistic ridge. The utility of griPEER is particularly apparent in cases with fully informative and largely informative 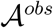; these cases correspond to low values of dissimilarity 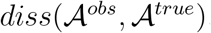 (marked at x-axis). As 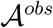 gets less informative about the true connections between coefficients in a model, MSEr of griPEER approaches MSEr of logistic ridge; these cases correspond to high values of dissimilarity 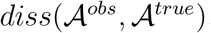. The result illustrates an important property of griPEER estimation method: adaptiveness to the amount of true information contained in an observed 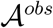 matrix. When 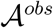 is largely informative, incorporating 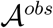 into the estimation is clearly a benefits. When 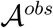 carries little or no information about the true connections between model coefficients, griPEER yields MSEr no larger than MSEr of logistic ridge estimator.

The performances of griPEER and logistic ridge depend on the structure of connections imposed by 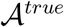 on the true *b*. We can observe that a difference between MSErs for griPEER and logistic ridge is smaller when 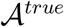 is defined based on 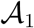: *homologous regions* matrix (Fig. 6, left column panel). Indeed, 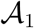 has smaller density than 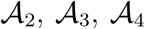 matrices, and imposes fewer connections between true coefficients in a model. Therefore, utilizing (full or partial) connectivity information 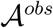 in estimation for 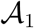-based signals is less beneficial compared to other considered patterns of coefficients dependencies. Furthermore, when each node is connected with everty other by a path consisting of strong connections, as in a case when 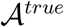 is created based on 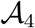 (4th column panel in Fig. 6), it is expected that all “true” model coefficients in a generated vector *b* are strongly dependent on each other; see Fig. 11 in Appendix B. In such a situation, even using even inaccurate information about the connections (high 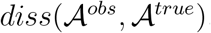 values) may be still be beneficial, as long as the correct message about strong coefficients’ dependence is provided. Finally, if we compare the results within each column panel of Fig. 6, we observe, as expected, that the estimation error gets smaller as number of predictors *p* gets smaller and as number of observations *n* gets larger.

**Figure 6:**
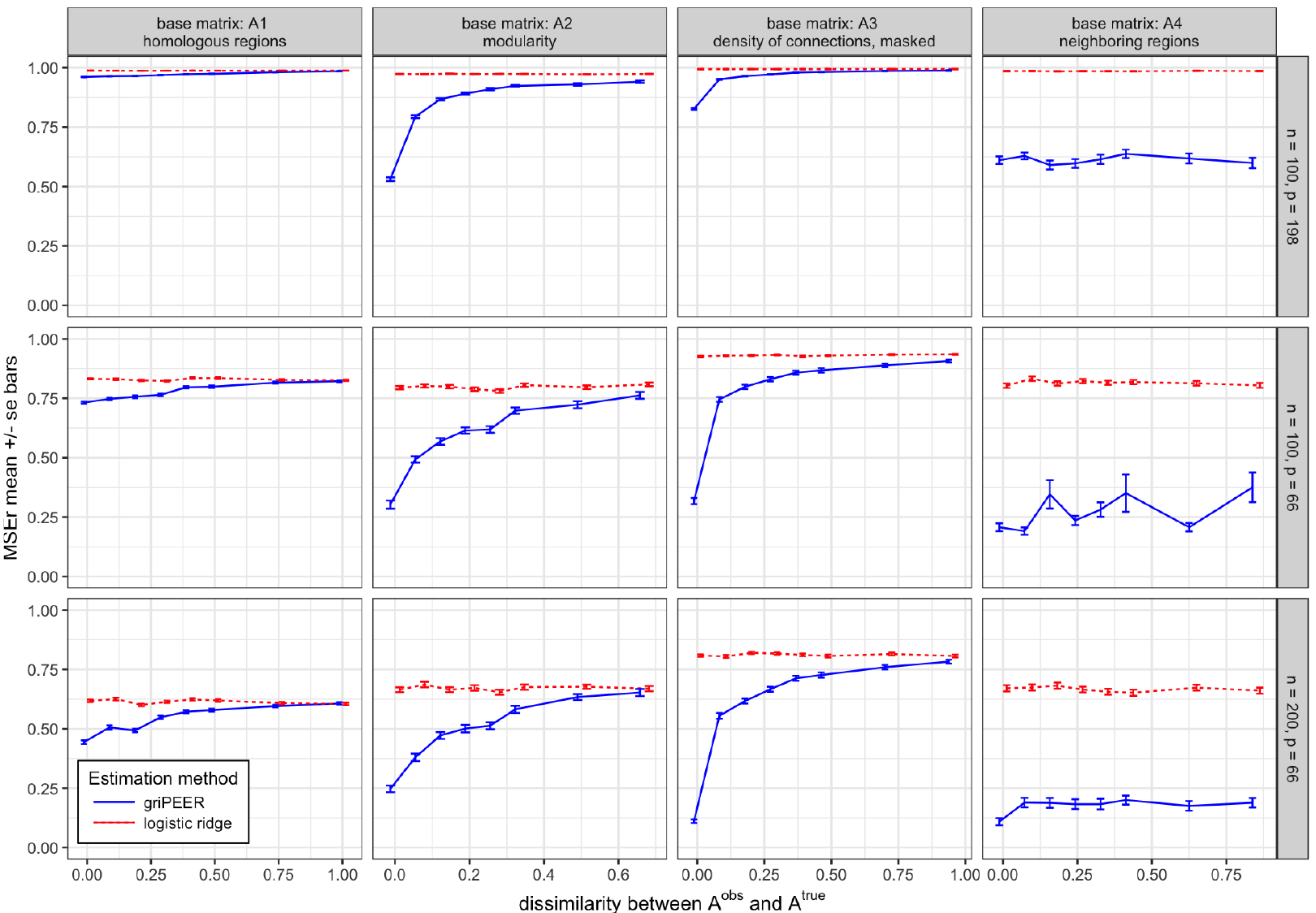
MSEr for estimation of *b* in Scenario 1. Results for griPEER (blue line) and logistic ridge (gray line). Presented are the average values of MSEr from 100 experiment runs for: *n* ∈ {100, 200}, *p* ∈ {66,198} and four true connectivity pattern inducing matrices, 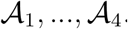. Dissimilarity between 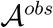 and 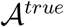 measured by 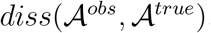 is represented by x-axis. Standard error of the mean bars are showed.

*Scenario 2*. In Scenario 2., we compare griPEER and logistic ridge estimation methods in a situation when an observed connectivity matrix, 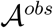, represents only positive connections, whereas 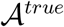 represents both positive and negative connections. We run the simulation for number of observations, *n* = 100, number of variables, *p* = 66, and for 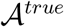 created based on four connectivity pattern inducing matrices, 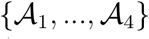. Matrix 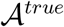 was generated form 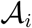 by switching signs in *k* columns (and corresponding rows), were *k* ∈ {1,4,7,10}. Fig. 7 displays the aggregated (mean) values of the relative estimation error based on 100 simulation runs.

With increasing *k*, 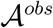 increasingly differs from the connectivity pattern used in the true signal generation and so the relative difference between MSEr for logistic ridge and griPEER decreases (for nearly all settings). Notably, MSEr for griPEER remains less than or equal to MSEr for logistic ridge. The results suggest that even using some incorrect information regarding the true connectivity structure (such as misspecifying negative dependencies as being positive) is not detrimental.

**Figure 7:**
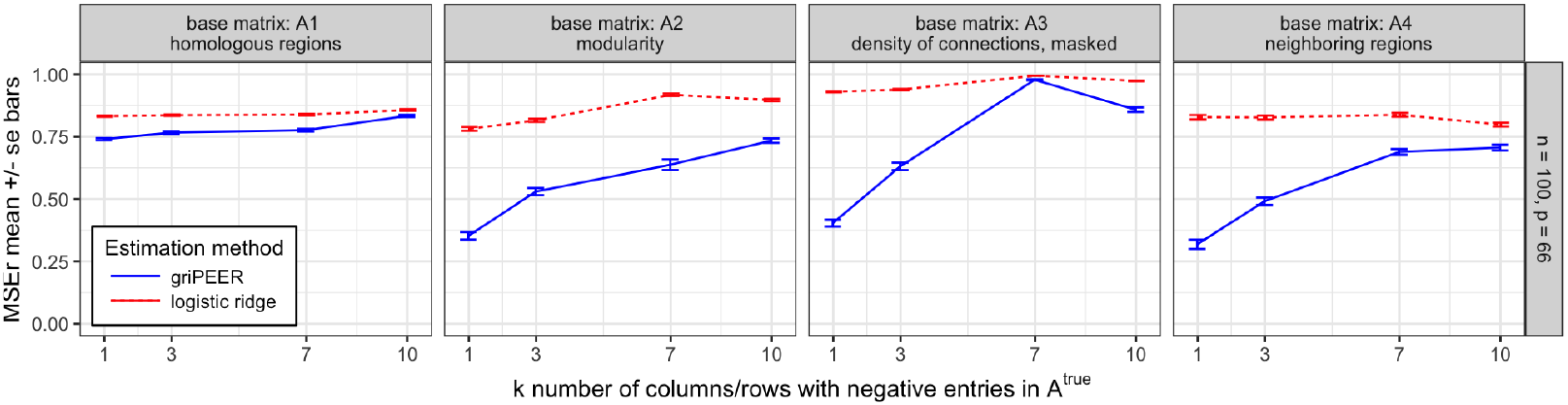
MSEr for estimation of *b* in Scenario 2. Results for griPEER (blue line) and logistic ridge (gray line). Presented are the average values of MSEr from 100 experiment runs for *n* = 100, *p* = 66 and four true connectivity pattern inducing matrices, 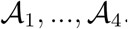. The number of columns (and corresponding rows) of 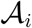, for which entries signs where switched in 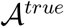 construction is represented by x-axis. Standard error of the mean bars are showed.

*Scenario 3*. In this scenario, we compare griPEER and logistic ridge estimation methods in a situation when 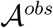 is of lower / higher matrix density than 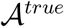. As in Scenario 2, we consider *n* = 100 and *p* = 66. This time, we do not change the signs of 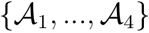 matrices but we generate 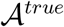 by adding/removing some connections to/from 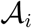. This influences the density of resulting matrix. In the simulation we consider 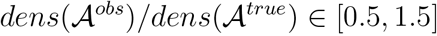 as a densities ratio range. Fig. 8 displays the mean values of the relative estimation error based on 100 simulation runs.

We can observe that, similar to Scenario 1, incorporating information on only a few connections (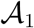 case) yields the smallest gain in the estimation accuracy measured by MSEr among all considered connectivity patterns. If 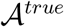 is set to 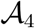, then (again, analogously to Scenario 1) the information about strong coefficients’ dependence is provided through 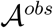. This results in substantially lower MSEr for griPEER across all densities ratio range we considered. When 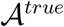 is equal to one of modules-based matrices, 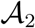 or 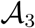, we still benefit from using 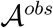 of lower density than 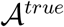, since 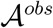 contains unaffected information about five separated modules in connectivity structure (values smaller than 1 at x-axis). Including the false connections in 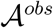 (values greater than 1 at x-axis) disturbs the message about the lack of dependencies between modules. A loss in griPEER’s estimation accuracy is apparent at the transition point *x* = 1. It remains, however, significantly better than the estimation accuracy for logistic ridge over the entire range of considered densities ratios.

**Figure 8:**
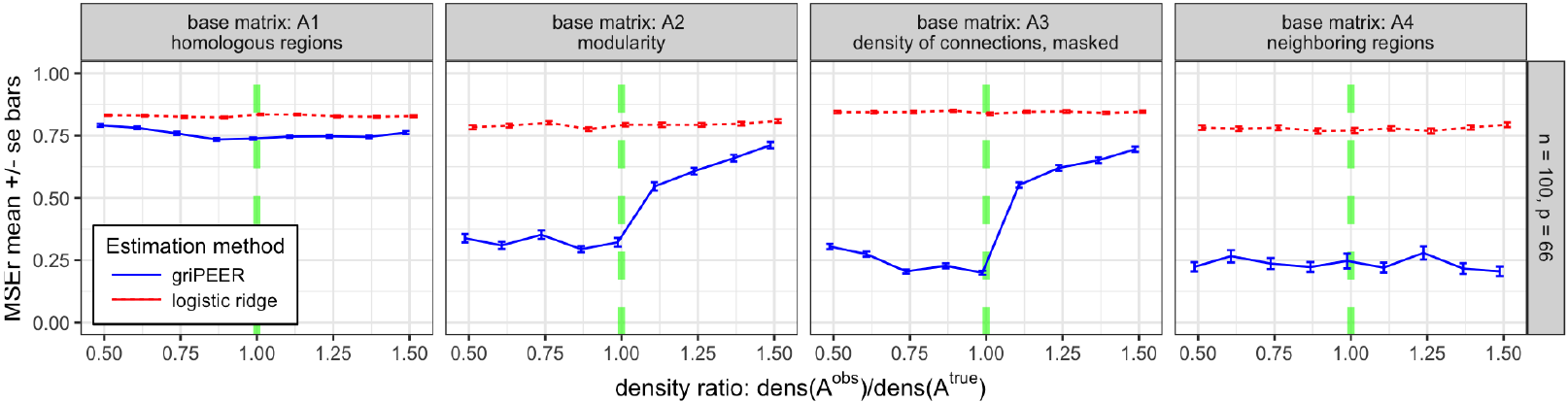
MSEr for estimation of *b* in Scenario 3. Results for griPEER (blue line) and logistic ridge (gray line). Presented are the average values of MSEr from 100 experiment runs for *n* = 100, *p* = 66 and four true connectivity pattern inducing matrices, 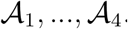. Ratio of densities, 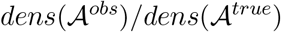, is represented by x-axis and varies from 0.5 to 1.5. Standard error of the mean bars are showed. Green dashed vertical lines denote the cases when ratio of matrix densities equals 1; in these cases, 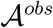 is identical to 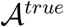.

### 5.3. Model coefficient significance testing

#### 5.3.1. Settings

We design a simulation study to evaluate performance of the two procedures for coefficient significance testing for griPEER, introduced in Section 4: asymptotic variance-covariance matrix-based approach, griPEER_asmp_, and Bootstrap-based approach, griPEER_boot_.

*Simulation scenario*. We follow the simulation setting used in Scenario 1, described in subsection 5.2. Specifically, we assume that 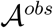 represents connections (partially) permuted with respect to connections represented by 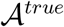. I.e., the corresponding 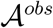 is constructed by randomizing entries in ***A***^*true*^ until a desired dissimilarity, 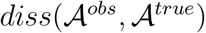, is achieved; see: Figure 3. The randomization technique preserves graph size, density, strength and graph degree-sequence (and hence degree distribution). Here, we confine ourselves to *p* = 66 and the case when 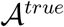 is based on 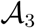struction of an adjacency matrix in the brain imaging analysis Section 4.

The adopted simulation scheme starts by generating the true signal and responses as in subsection 5.2. We generate large number of observations, *n* = 1000, but in the estimation we use only 150 records to emulate a real data setting. The large sample size is used only to label the variables which are “truly relevant” so that the performance of griPEER_asmp_ and griPEER_boot_ in the context of variables selection can be assessed. Defining “truly relevant” variables is done through the asymptotic confidence interval for the logistic model estimate (non-regularized estimation), which is unbiased and asymptotic normal (Fahrmeir and Kaufmann, 1985). The details are described below.

*Simulation study procedure*. In the experiment, we perform the following steps.

1. Apply steps 1-5 from the simulation study procedure described in subsection 5.2 with *n* = 1000.
2. Run the following steps 100 times:

a. generate *p*-dimensional vector of true coefficients, *b*, as well as *n*-dimensional vectors, *θ* and *y*, by following steps 2(a)–2(d) described in subsection 5.2,
b. calculate the asymptotic standard deviations, 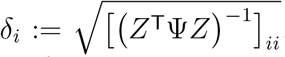, for *i* = 1,…, *p*, where 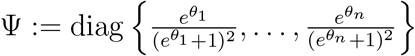 (see, Appendix A.3),
c. divide the set of indices, {1,…, *p*}, into two separated groups: *I*_*T*_, corresponding to the variables defined as *relevant* and *I*_*F*_, corresponding to the variables defined as *irrelevant*, by using the criterion

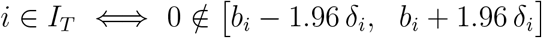
d. generate the data for estimation, *y*\*, *X*\* and *Z*\*, by taking first 150 rows of *y*, *X* and *Z*; center and normalize the columns of *Z*\* to zero means and unit ℓ_2_ norms,
e. apply griPEER_asmp_ and griPEER_boot_ on *y*\*, *X*\* and *Z*\* to indicate response-related variables defined by each of methods,
f. based on information about “truly relevant” and “truly irrelevant” variables; i.e., the known division into *I*_*T*_ and *I*_*F*_, for each method identify: *S* — the number of true discoveries and *V* — the number of false discoveries,
g. for each method collect measures 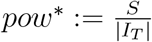 and 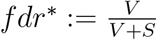,
3. Define the estimates of *power* and *FDR* as the averages of *pow*\* and *fdr*\* (across 100 repetitions of the step 2).

#### 5.3.2. Results

Figure 9 displays the values of power (left plot) and FDR (right plot), estimated based on the simulation procedure described in subsection 5.3.1. As expected, for both methods power decreases as 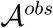 becomes less informative regarding the true connections between coefficients in a model. We observe however, that griPEER_boot_ is able to reach substantially higher power than griPEER_asmp_ under considered settings. The estimated FDRs are not very distinct for both methods and they tend to be very similar for less accurate connectivity information.

The results obtained in the simulation study suggest that one can potentially gain power by utilizing griPEER_boot_ for coefficient significance testing, compared to griPEER_asmp_ approach. In addition, power gain occurs without a substantial increase of FDR. consequently, we employ griPEER_boot_ in the real data application in Section 6.

**Figure 9:**
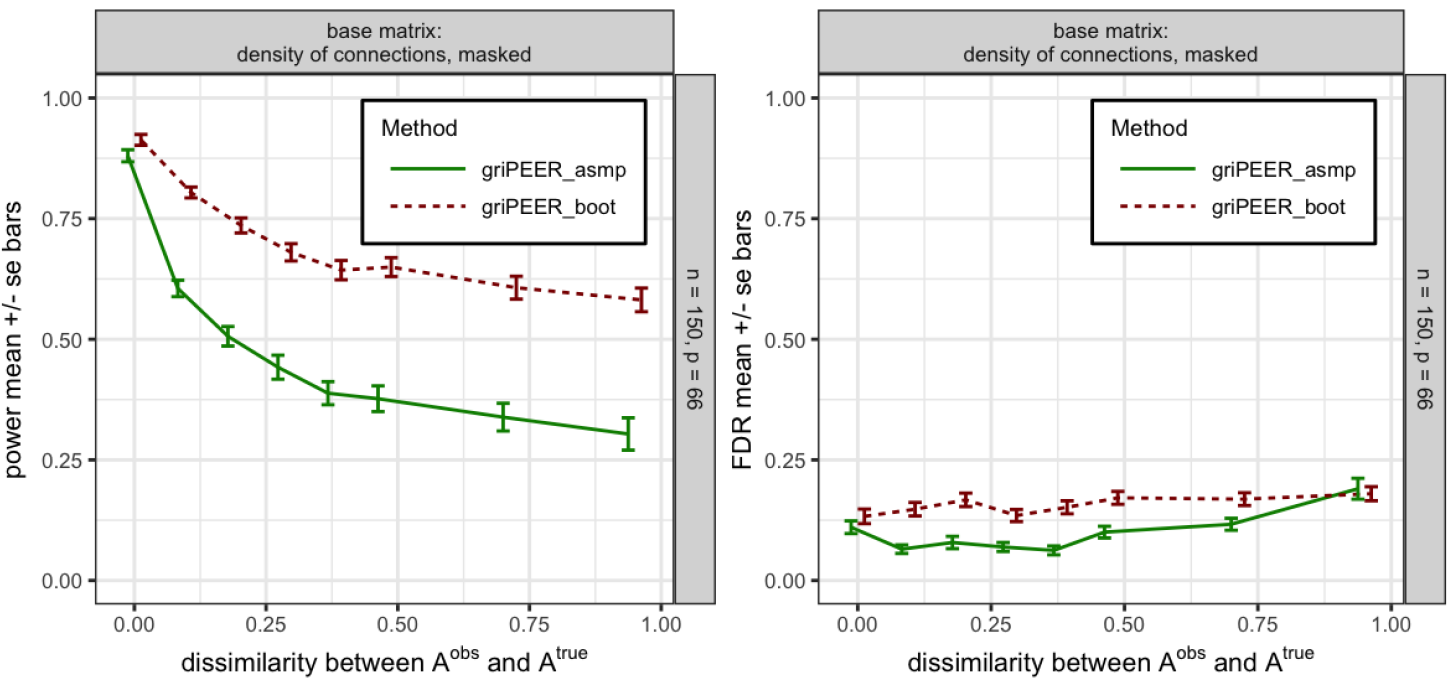
The estimated values for power (left plot) and FDR (right plot) obtained with asymptotic variance-covariance matrix-based approach (griPEER_asmp_, blue line) and Bootstrap-based approach (griPEER_boot_; gray line). Values are aggregated (mean) out of 100 experiment runs, for number of observations *n* = 150, number of variables *p* = 66 and 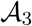 (Figure 2, middle right plot) as a true connectivity pattern. The dissimilarity between 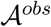 and 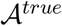 is represented by the x-axis. Standard error of the mean bars are showed.

### 5.4. The software used in simulations

The code used to generate the results was built in Matlab and is available at GitHub (https://github.com/dbrzyski/griPEER).

## 6. Imaging data application

We model the association between the presence/absence of HIV and the properties of the structural cortical brain imaging data. More specifically, we employ cortical thickness measurements obtained using the FreeSurfer software (Fischl, 2012) to classify the binary response indicating the status of HIV infection, where 0 indicates an HIV-negative individual and 1 an HIV-positive individual.

### 6.1. Data and preprocessing

*Study sample*. The analyzed sample consists of 162 young (age range: 18-42 years) males, where 108 were HIV-positive and 54 were HIV-negative. Study sample subjects’ demographic and a HIV-related characteristics are summarized in Table 1.

**Table 1:**
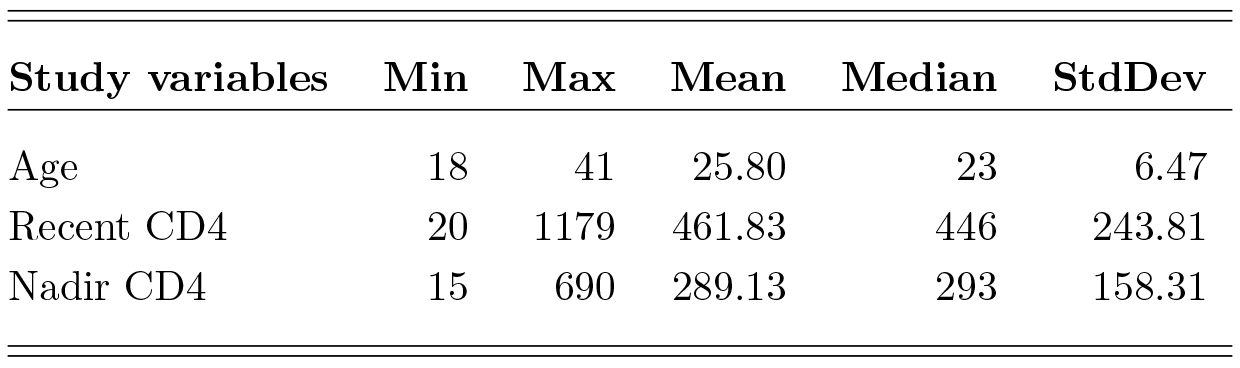
Study sample subjects’ characteristics.

| Study variables | Min | Max | Mean | Median | StdDev |
| --- | --- | --- | --- | --- | --- |
| Age | 18 | 41 | 25.80 | 23 | 6.47 |
| Recent CD4 | 20 | 1179 | 461.83 | 446 | 243.81 |
| Nadir CD4 | 15 | 690 | 289.13 | 293 | 158.31 |

*Cortical measurements*. The FreeSurfer software package (version 5.1) was used to process the acquired structural MRI data, including gray-white matter segmentation, reconstruction of cortical surface models, labeling of regions on the cortical surface and analysis of group morphometry differences. The resulting dataset has cortical measurements for 68 cortical regions with parcellation based on Desikan-Killiany atlas (Desikan et al., 2006). The subset of 66 variables describing average gray matter thickness (in millimeters) of gray matter brain regions did not incorporate left and right insula due to their exclusion from the structural connectivity matrix.

*Structural connectivity information*. In the analysis we used two adjacency matrices, which were incorporated in the estimation with griPEER through the normalized Laplacian matrix. The adjacency matrices were created based on two structural connectivity information types: density of connections (DC) and fractional anisotropy (FA). For each of them, two steps were performed to achieve the final adjacency matrix, 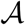. In the first step, we computed the entry-wise median (across subjects) of DC or FA connectivity matrices. The second step relied on “masking by modularity partition”, i.e. limiting the information achieved in the first step only to the connections between brain regions being in the same modules (i.e. we set 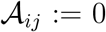, if regions *i* and *j* were not in the same module). For this purpose, we used the modularity connectivity matrix (see Sporns (2013); Cole et al. (2014); Sporns and Betzel (2016)), which defines the division of the brain into five separated communities. The modularity matrix was obtained by using Louvain method (Blondel et al., 2008) and based on model proposed in Hagmann et al. (2008). More details on this construction can be found in Karas et al. (2017).

### 6.2. Estimation methods

We employed logistic ridge and griPEER_boot_ to classify the HIV-infected and non-infected individuals based on the estimate cortical thickness measurements. All analyses were adjusted for *Age* with its respective coefficient non-penalized. Consequently, *X* was an *n* by 2 matrix containing the column of ones (representing the intercept) and the column corresponding to subjects’ age. Columns of design matrices (other than intercept) were centered to zero mean and normalized to unit standard deviation before the estimation. The selection of regularization parameter in logistic ridge was done within the GLMM framework. For all methods we used Bootstrap-based approach with 50,000 samples, to define the subset of statistically significant variables.

### 6.3. Results

The estimates obtained from the logistic ridge and the griPEER_boot_ for considered groups of subjects are presented in Figure 10. Brain regions labeled as response-related are marked with solid red vertical lines. In Table 2, we summarize the estimated values corresponding to brain regions being labeled as response-related by at least one considered approach. Note that all significant associations are negative, indicating thinner cortical areas are indicative of HIV-positive status. Significant estimates obtained from the griPEER for both types of connectivity matrices (FA- and DC-based) agree in 7 out of 8 cortical brain regions, while the logistic ridge significant findings disagree with the FA-based griPEER estimates in 4 regions and with the DC-based griPEER estimates in 3 regions.

**Figure 10:**
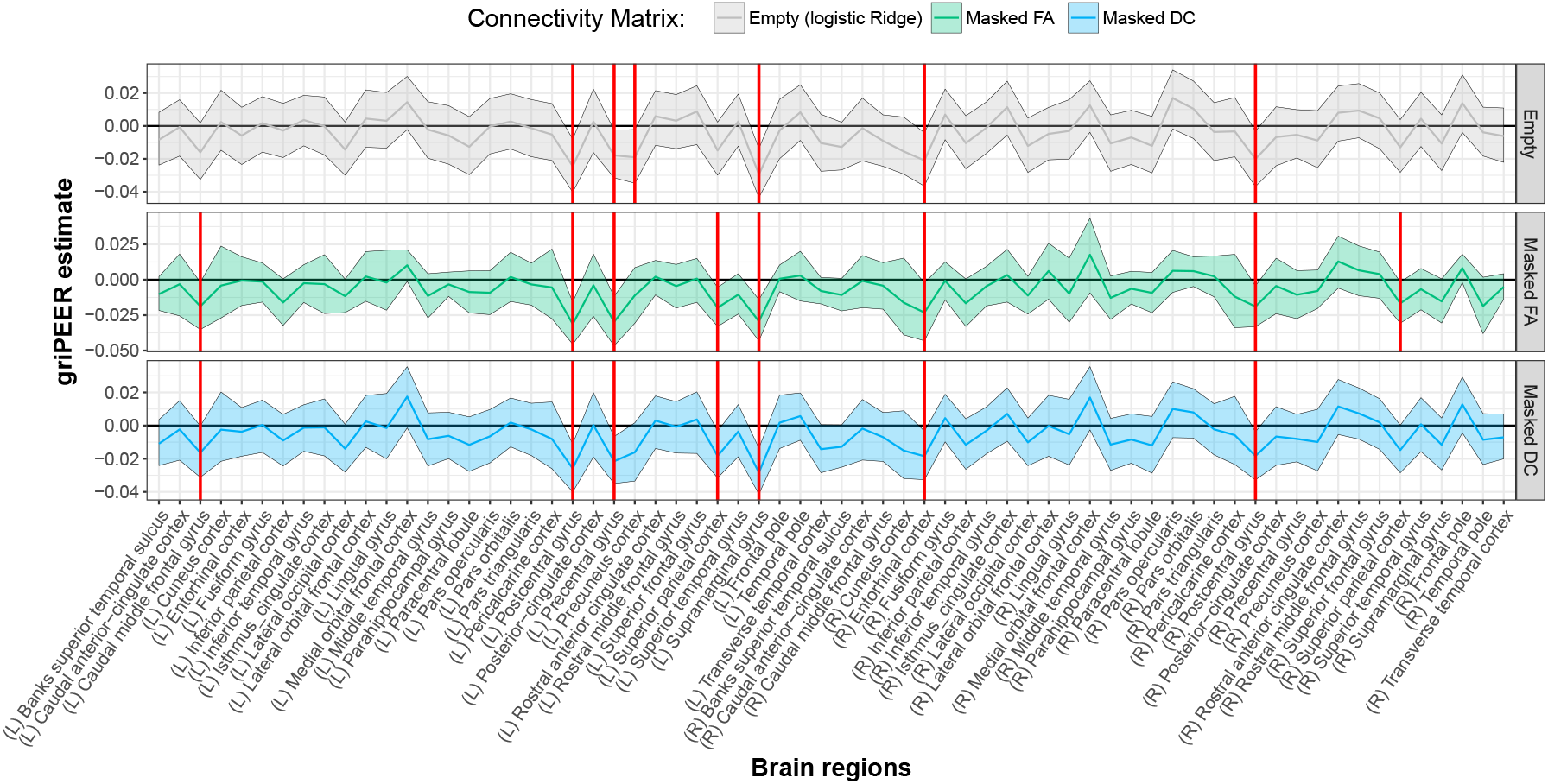
Results obtained by griPEER_boot_ on 162 subjects (with 108 HIV-infected). Here, the response variable was defined as the disease indicator and 66 cortical brain regions were considered - 33 from the left and 33 from the right hemisphere. Regions labeled as response-related were marked by red vertical lines. Confidence intervals were calculated based on 50,000 bootstrap samples.

## 7. Discussion

We have provided a rigorous and computationally feasible method which incorporates additional information to estimate regression parameters in the generalized linear model setting. Our proposed method, griPEER, extends our work performed in the linear model setting Karas et al. (2017). We utilize the structural connectivity information obtained from the DTI to inform the association between the cortical covariates and a generalized outcome (e.g. binary indicator of HIV-infection). The structural connectivity information is used to create a Laplacian matrix, which in turn is used to specify the regularization penalty.

**Table 2:** Estimates of the cortical brain regions coefficients obtained by the logistic ridge and griPEER_boot_ with two different connectivity matrices – fractional anisotropy (Masked FA) and density of connections (Masked DC). Both matrices were masked by the modularity matrix before the analysis. Values corresponding to regions being labeled as response-related are shown in bold, green font and non-significant findings are shown using the red font. We show the results for all regions being selected by at least one method as response-related.

| Connectivity type | Caudal middle frontal [L] | Post central [L] | Pre central [L] | Pre cuneus [L] | Superior parietal [L] | Supra marginal [L] | Entorhinal [R] | Post central [R] | Superior parietal [R] |
| --- | --- | --- | --- | --- | --- | --- | --- | --- | --- |
| Empty | **-0.016** | **-0.025** | **-0.018** | **-0.019** | **-0.015** | **-0.029** | **-0.021** | **-0.020** | **-0.013** |
| Masked FA | **-0.019** | **-0.031** | **-0.030** | **-0.011** | **-0.020** | **-0.029** | **-0.023** | **-0.019** | **-0.017** |
| Masked DC | **-0.016** | **-0.026** | **-0.022** | **-0.016** | **-0.018** | **-0.028** | **-0.018** | **-0.018** | **-0.015** |

The simulation study shows that in each scenario considered, the proposed method, griPEER, outperforms logistic ridge in a binomial model coefficient estimation - griPEER yields smaller or similar estimation relative error 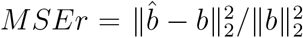 compared to the logistic ridge. Performance of griPEER is significantly better when the observed connectivity information is fully or largely informative about the true connectivity structure between model coefficients. Notably, even in cases when observed connectivity information is only partially informative or completely non-informative, the proposed method yields MSEr no larger than the logistic ridge estimator.

Application of griPEER to classify the individuals as HIV-infected and noninfected resulted in discovery of 3 additional cortical regions, namely Left Caudal Middle Frontal Gyrus, Left Superior Parietal Lobule and Right Superior Parietal Lobule, that were thinner in the HIV-infected individuals.

Our future work will incorporate both structural and functional brain connectivity information in the regularized estimation procedure. We will also include other properties of the cortex, namely the cortical area and its curvature.

## Acknowledgements

Research support was partially supported by the NIMH grants R01MH108467.

## A. Appendix

## A.1. Proof of proposition 3.1

The claim quickly follows from two well known linear algebra theorems: Woodbury identity and matrix determinant lemma. We recall the both results below as Theorem A.1.

### Theorem A.1.

*Suppose that A and C are invertible n by n matrices and U, V are n by p matrices. Then*

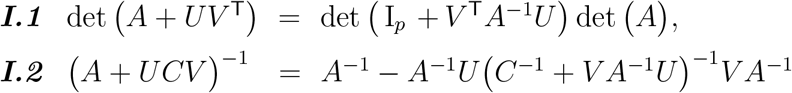

We will start with the proof of (C.1). Thanks to I.1,

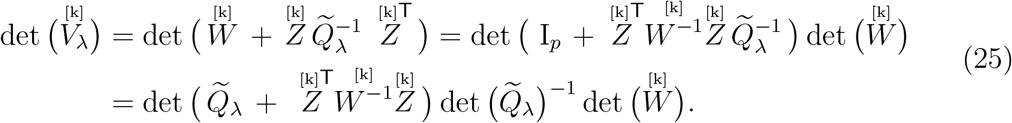

Now

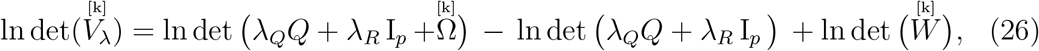

which finishes the proof (C.1). To show the second claim, we will rewrite 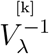 as

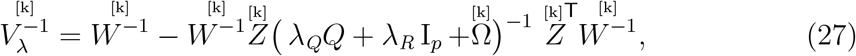

thanks to (I.2). Therefore

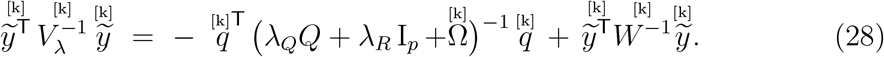

## A.2. Gradient and Hessian for the objective in (18)

Denote by *h*(λ_*Q*_, λ_*R*_) the objective function of interest, i.e.

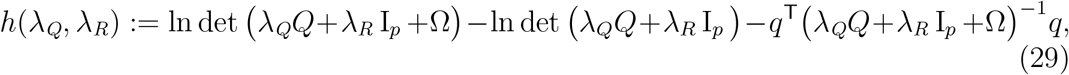

where Ω and *q* were defined in the statement of proposition 3.1 (“[k]”s symbols were omitted for clarity). After using notations *D*_λ_ := (λ_*Q*_*Q* + λ_*R*_ I_*p*_ +Ω)^−1^ and 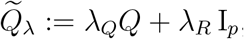, this function takes the short form

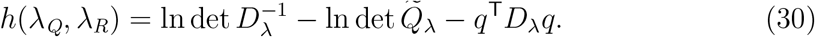

To find the gradient and Hessian of *h* we will use the following well known formulas

### Proposition A.2.

*Suppose that A and B are p by p, symmetric, positive semi-definite matrices, ν is p is p-dimensional vector and tA + sB is positive definite*.

*Then it holds*

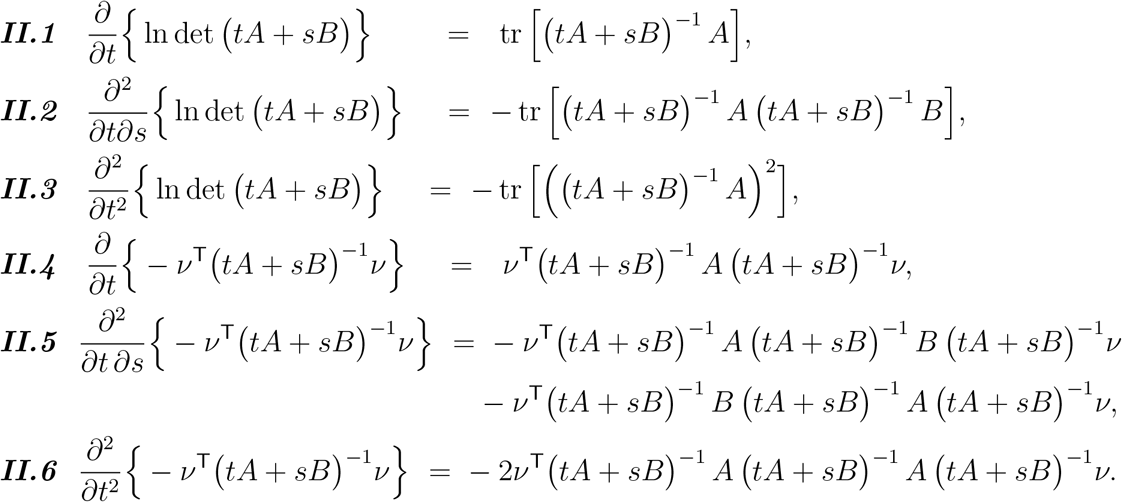

Thanks to the above, we quickly get

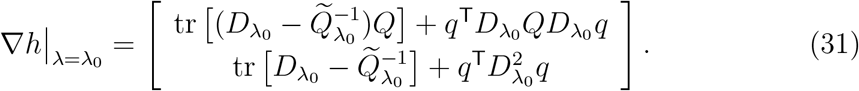

and

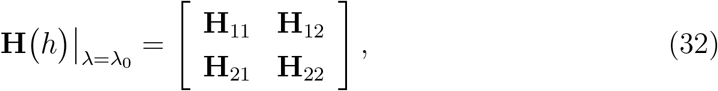

where

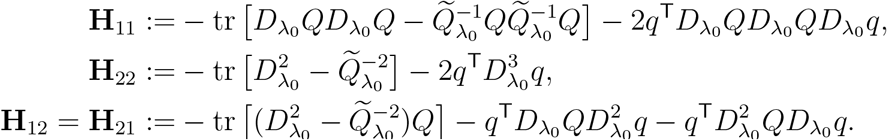

## A.3. Asymptotic confidence interval

We start with the optimization problem equivalent to 20, with the objective multiplied by 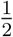,

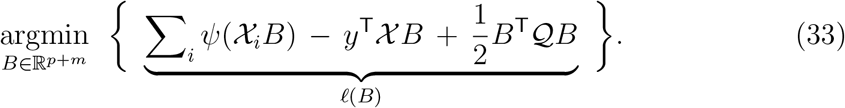

Calculating the derivatives of *ℓ* yields

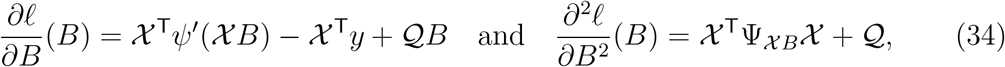

where

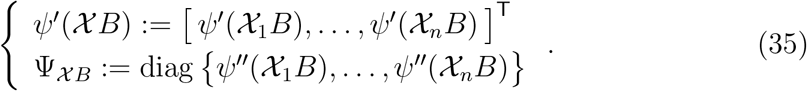

Denote by *B*_T_ the true signal and consider the Taylor series expansion of 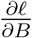 about *B*_T_. If we consider the value of Taylor polynomial in the solution of (33), 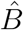, this yields the following expression

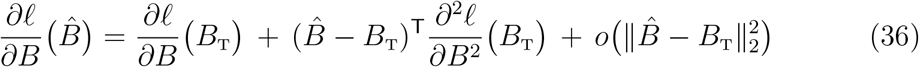

Since the left-hand side of the above equals zero, using (34) we get the first-order approximation of 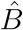

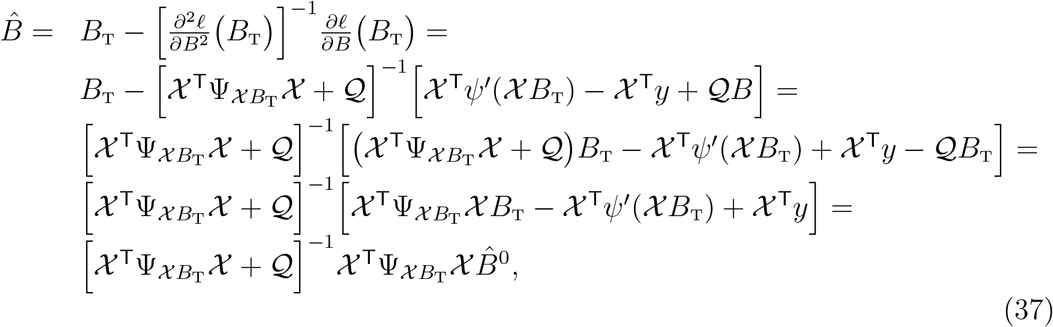

where 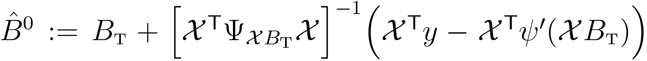 is the first-order approximation of the generalized linear model estimate, i.e. for 𝒬 = 0. It was shown that, under some regularity conditions, this estimate is unbiased and asymptotic normal (Fahrmeir and Kaufmann, 1985). The corresponding asymptotic variance is 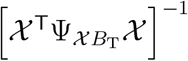. Consequently, the asymptotic variance, *var*_*a*_, of 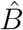 is given by

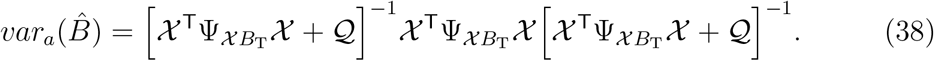

## B. Appendix

**Figure 11:**
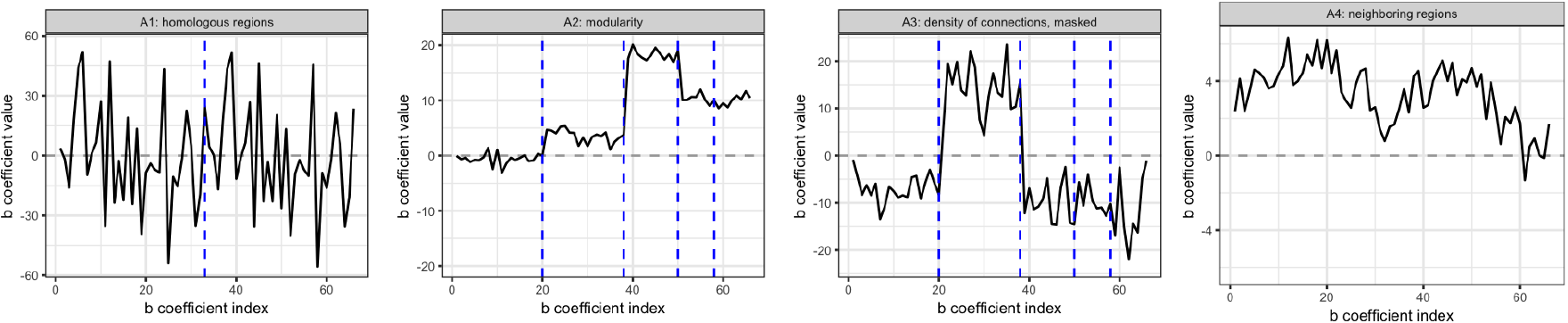
Examplary vectors of *b* model coefficients generated as 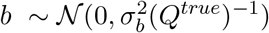, where *Q*^*true*^ is Laplacian matrix of 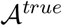 graph adjacency matrix. Clearly, *b* coefficient values reflect the connectivity structure represented by 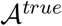 matrices assumed in the simulation study; left plot: 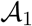 “*homologous regions*”, middle left plot: 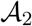 *“modularity”*, middle right plot: 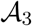 *“density of connections, masked”*, right plot: 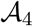 *“neighboring regions”* (see: Fig. 2). In the left plot, vertical dashed line marks the separation between coefficients corresponding to left hemisphere brain regions and right remishpere brain regions assumed in 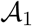 *“homologous regions”* construction. In the middle left plot, vertical dashed lines mark the separation between connectivity modules assumed in 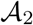 *“homologous regions”* construction. In the middle right plot, vertical dashed lines mark the separation between connectivity modules assumed in 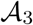 *“homologous regions”* construction.

## References

Bertero, M., Boccacci, P., 1998. Introduction to Inverse Problems in Imaging. Institute of Physics, Bristol, UK.

Blondel, V. D., Guillaume, J. L., Lambiotte, R., Lefebvre, E., 2008. Fast unfolding of communities in large networks. Journal of statistical mechanics: theory and experiment 2008 (10).

Breslow, N. E., Clayton, D. G., 1993. Approximate inference in generalized linear mixed models. Journal of the American Statistical Association 88 (421), 9–25.

Cessie, S. L., Houwelingen, J. C. V., 1992. Ridge estimators in logistic regression. Journal of the Royal Statistical Society. Series C (Applied Statistics) 41 (1), 191–201.

Chung, F., 2005. Laplacians and the cheeger inequality for directed graphs. Annals of Combinatorics 9 (1), 1–19.

Cole, M. W., Bassett, D. S., Power, J. D., Braver, T. S., Petersen, S. E., 2014. Intrinsic and task-evoked network architectures of the human brain. Neuron 83 (1), 238–251.

Desikan, R. S., Segonne, F., Fischl, B., Quinn, B. T., Dickerson, B. C., Blacker, D., Buckner, R. L., Dale, A. M., Maguire, R. P., Hyman, B. T., Albert, M. S., Killiany, R. J., 2006. An automated labeling system for subdividing the human cerebral cortex on mri scans into gyral based regions of interest. NeuroImage 31 (3), 968–80.

Engl, H. W., Hanke, M., Neubauer, A., 2000. Regularization of inverse problems. Kluwer, Dordrecht, Germany.

Fahrmeir, L., Kaufmann, H., 1985. Consistency and asymptotic normality of the maximum likelihood estimator in generalized linear models. Ann. Statist. 13 (1), 342–368.

Fischl, B., Aug. 2012. FreeSurfer. Neuroimage 62 (2), 774–81. URL http://www.ncbi.nlm.nih.gov/pmc/articles/PMC3685476/

Hagmann, P., Cammoun, L., Gigandet, X., Meuli, R., Honey, C. J., Wedeen, V. J., Sporns, O., 2008. Mapping the structural core of human cerebral cortex. PLoS Biol 6 (7), e159.

Hastie, T., Buja, A., Tibshirani, R., 1995. Penalized discriminant analysis. The Annals of Statistics 23 (1), 73–102.

Huang, J., Shen, H., Buja, A., 2008. Functional principal components analysis via penalized rank one approximation. Electronic Journal of Statistics 2, 678–695.

Karas, M., Brzyski, D., Dzemidzic, M., Goni, J., Kareken, D. A., Randolph, T. W., Harezlak, J., 2017. Brain connectivity-informed regularization methods for regression. Preprint available on the bioXiv:10.1101/117945.

Li, C., Li, H., 2008. Network-constrained regularization and variable selection for analysis of genomic data. Bioinformatics 24 (9), 1175–1182.

Maldonado, Y. M., 2009. Mixed models, posterior means and penalized least-squares. Optimality 57, 216–236.

Phillips, D., 1962. A technique for the numerical solution of certain integral equations of the first kind. Journal of the ACM 9 (1), 84–97.

Pinheiro, J. C., Chao, E. C., 2006. Efficient laplacian and adaptive gaussian quadrature algorithms for multilevel generalized linear mixed models. Journal of Computational and Graphical Statistics 15 (1), 58–81.

Randolph, T. W., Harezlak, J., Feng, Z., 2012. Structured penalties for functional linear models – partially empirical eigenvectors for regression. Electronic Journal of Statistics 6, 323–353.

Slawski, M., Castell, W. Z., Tutz, G., 2010. Feature selection guided by structural information. Annals of Applied Statistics 4 (2), 1056–1080.

Sporns, O., 2013. Network attributes for segregation and integration in the human brain. Current opinion in neurobiology 23 (2), 162–171.

Sporns, O., Betzel, R. F., 2016. Modular brain networks. Annual review of psychology 67, 613.

Tibshirani, R., 1996. Regression shrinkage and selection via the lasso. Journal of the Royal Statistical Society: Series B 58 (1), 267–288.

Tibshirani, R., Saunders, M., Rosset, S., Zhu, J., Knight, K., 2005. Sparsity and smoothness via the fused lasso. Journal of the Royal Statistical Society: Series B 67 (1), 91–108.

Tibshirani, R., Taylor, J., 2011. The solution path of the generalized lasso. The Annals of Statistics 39 (3), 1335–1371.

Tikhonov, A., 1963. Solution of incorrectly formulated problems and the regularization method. Soviet Math 4 (4), 1035–1038.

Wolfinger, R., O’connell, M., 1993. Generalized linear mixed models a pseudolikelihood approach. Journal of Statistical Computation and Simulation 48 (3), 233–243.

Xin, B., Kawahara, Y., Wang, Y., Gao, W., 2016. Efficient generalized fused lasso and its applications. ACM Transactions on Intelligent Systems and Technology (TIST) 7 (4).

Zeger, S., Karim, M. R., 1991. generalized linear models with random effects - a gibbs sampling approach. Journal of the American Statistical Association 86 (413), 79–86.

Zhao, S., Shojaie, A., 2016. A significance test for graph-constrained estimations. Biometrics 72 (2), 484–493.

Zou, H., 2006. The adaptive lasso and its oracle properties. Journal of the American Statistical Association 101 (476), 1418–1429.

Zou, H., Hastie, T., 2005. Regularization and variable selection via the elastic net. Journal of the Royal Statistical Society: Series B 67 (2), 301–320.

